# An integrative taxonomic approach reveals unexplored diversity in Croatian planarians

**DOI:** 10.1101/2025.04.10.648211

**Authors:** Miquel Vila-Farré, Jeremias N. Brand, Tobias Boothe, Maren Brockmeyer, Fruzsina Ficze-Schmidt, Markus A. Grohme, Uri Weill, Kasper H. Kluiver, Ludwik Gąsiorowski, Lucija Kauf, Yuliia Kanana, Helena Bilandžija, Marta Riutort, Jochen C. Rink

**Affiliations:** Department of Tissue Dynamics and Regeneration, Max Planck Institute for Multidisciplinary Sciences, Göttingen, Germany; Max Planck Institute of Molecular Cell Biology and Genetics, Dresden, Germany; Computing, Mathematics, Engineering & Natural Sciences Faculty, Northeastern University London, London, United Kingdom; Institute of Evolutionary Biology, Faculty of Biology, University of Warsaw, Warsaw, Poland; Ruđer Bošković Insititute, Bijenička cesta 54, Zagreb, Croatia; Institute of Neurogenetics, University of Lübeck, Lübeck, Germany; I.I. Schmalhausen Institute of Zoology of National Academy of Sciences of Ukraine, Kyiv, Ukraine; Departament de Genètica, Microbiologia i Estadística, Universitat de Barcelona, Barcelona, Spain; Institut de Recerca de la Biodiversitat (IRBio), Universitat de Barcelona, Barcelona, Spain; Faculty of Biology and Psychology, Georg-August-University Göttingen, Göttingen, Germany

**Keywords:** diversity, barcoding, *cytochrome c oxidase subunit I*, taxonomy, planarians, triclads, Croatia, Western Balkans

## Abstract

**Background:** Freshwater ecosystems are among the most endangered habitats on Earth, with approximately one-fourth of aquatic species at risk of extinction. Effective conservation efforts require comprehensive monitoring and accurate species identification, including often overlooked groups. DNA barcoding promises rapid and accessible species identification but requires the availability of “universal” primer pairs and robust, taxonomically curated reference libraries. Planarian flatworms are one such group for which these resources are currently lacking, even though they are common constituents of freshwater ecosystems worldwide. As a result, the true extent of planarian diversity remains underappreciated in many areas.

**Results:** Motivated by the highly skewed representation of planarian diversity in current GenBank records of the barcoding gene mitochondrial *cytochrome c oxidase subunit I* (COI), we optimised existing barcoding primers and tested them on planarians collected in a species-rich but, so far, poorly surveyed region of Croatia. Using an integrative approach combining the new primers, traditional taxonomy and karyology, we generated new taxonomically curated COI barcode sequences for European dugesiids, dendrocoelids and planariids, which, in the case of the latter two groups, represent a substantial increase of the number of entries for those families. Our efforts resulted in several significant findings, including the description of a new pigmented *Dendrocoelum* species, *Dendrocoelum pigmentatum* sp. nov, the discovery of two highly differentiated haplotypic clades in *Schmidtea lugubris*, and the rediscovery of *Polycladodes alba* in Croatia after a century.

**Conclusions:** Overall, our work extends the utility of DNA barcoding for species-level identification of planarians to previously inaccessible groups. Additionally, it integrates Croatia as an underexplored but species-rich region into the endeavour to systematically describe the planarian fauna of Europe. The expansion of a known number of Croatian planarian species from eight to sixteen and the discovery of a new, large planarian species in continental Europe, *Dendrocoelum pigmentatum*, demonstrate the effectiveness of our integrative approach. Overall, our work highlights the underappreciated diversity of planarians, even in continental Europe and supports practical conservation efforts to preserve aquatic biodiversity.

## Introduction

Assessing organismal diversity is critical amidst the ongoing biodiversity crisis that threatens global ecosystems(1, 2). The decline in vertebrate biodiversity, such as amphibians, is well-documented(3). In contrast, invertebrates, which contain 99 % of the animal diversity(4), remain, with few exceptions(5), understudied. As a result, only a small proportion of invertebrates are considered for conservation actions(6). This knowledge gap is particularly concerning for freshwater ecosystems, which are among Earth’s most endangered habitats(7). Freshwater ecosystems account for the highest proportion of species extinctions and harbour many threatened species, with approximately one-fourth currently at risk of extinction(8). This figure likely underestimates the crisis’ true extent, as many data-deficient groups remain poorly studied(9, 10), underscoring the need for increased research and monitoring(11). Accurate species identification and description are essential for determining priorities for biomonitoring and conservation actions(11).

However, species identification via traditional taxonomy is often time-intensive and demands specialised expertise. DNA barcoding has emerged as a powerful tool to address these challenges(12). The term refers to the sequencing of DNA regions that evolve so rapidly that they tend to accumulate species- or even population-specific single-nucleotide polymorphisms (SNPs). Particularly useful loci are additionally flanked by slowly evolving regions to allow PCR amplification of the intervening hypervariable region by common primer sets in a broad range of species. Commonly used barcoding loci include the nuclear ribosomal rRNA genes (e.g., 18S) for genus or species-level identification and the mitochondrial *cytochrome c oxidase subunit I* (COI) locus for species or population-level analyses(13, 14). However, effective DNA barcoding depends on the availability of “universal” primer pairs that robustly amplify the barcoding locus across the target taxon and taxonomically curated reference libraries for barcode-based species identification—resources often lacking for many invertebrate groups(15–20).

Planarians (Order Tricladida) are renowned for their remarkable regenerative capabilities, making species like *Schmidtea mediterranea* and *Dugesia japonica* models in stem cell and regeneration research(21–23). Recent studies have emphasised the need to incorporate additional planarian species to better understand the evolutionary processes underlying regeneration(24–27). However, the actual diversity of planarians is poorly understood. This is more striking given their worldwide distribution, abundance, and potential for studying a wide range of biological questions(23, 28–32). Consequently, the conservation status of many planarians is unknown. The decline of the model species *S. mediterranea* in Spain highlights the need for efforts in planarian conservation(33). Despite this, the IUCN Red List, a recognised international reference inventory of biological species’ global conservation status and extinction risk(34), lists only two planarian species as assessed in 2024(10). One of the reasons is that traditional taxonomy in planarians is carried out via the reconstruction of copulatory apparatus morphology from histological sections, which is time-consuming and requires specialist training and extensive experience(35); on the other hand, barcoding species delimitation has been primarily used in dugesiids of the Mediterranean area and land planarians, but rarely so far in other planarian groups(36–40).

Croatia lies within the Mediterranean biodiversity hotspot(41) and at the intersection of several biogeographic regions(42), with a dynamic geological history contributing to its rich biodiversity. Our review of taxonomically curated records identified at least eight freshwater planarian species in the country, primarily based on studies from the 20th century(43–51): *Polycelis felina* (Dalyell, 1814), *Phagocata dalmatica* (Stanković and Komárek, 1927), *Crenobia alpina* (Dana, 1766), *Crenobia anophtalma* Mrázek, 1907, *Dendrocoelum romanodanubiale* (Codreanu, 1949), *Dendrocoelum subterraneum* Komárek, 1919, *Polycladodes alba* Steinmann, 1910, *Dugesia absoloni* (Komárek,1919). Records of planarian species that are not taxonomically documented are not included in this list (e.g., (52, 53)). This relatively low species count suggests that Croatia’s freshwater planarian diversity is likely underestimated.

To address this potential gap, we surveyed freshwater planarians in Croatia’s karstic region. Our study integrates DNA barcoding using newly designed freshwater planarian primers, RNA sequencing, and traditional taxonomic methods, including histological analyses and metaphase chromosome preparations. Our research expands the number of known Croatian planarian species from eight to sixteen, including the discovery and description of a new, large-pigmented dendrocoelid species. In addition, we significantly advance the utility of barcoding for planarian taxonomy by developing new primers and targeting previously unstudied planarian clades, thus facilitating species identification and monitoring in this understudied group.

## Methods

### Field collections

The specimens analysed in this study were collected via dedicated field expeditions to Croatia and other parts of Europe, mostly between 2013 and 2023. Planarians were sampled from the underside of stones, aquatic plants or other submerged objects with the help of a soft brush to minimise damage to the animals(27). After collection, specimens were transferred into 50 ml conical bottom tubes, with care taken to maintain low animal densities to prevent worm lysis. The tubes were kept cool, and water from the collection site was used for daily water changes during the field campaign to maintain health and viability. Each collection site’s GPS coordinates and detailed habitat characteristics were recorded in a dedicated database for future reference.

### Live imaging

Flatworms were imaged either in the field or under laboratory conditions. Field imaging was conducted using a Canon EOS 6D Mark II camera equipped with a Canon EF 100mm f/2.8L Macro IS USM lens and a Canon Extension Tube EF25 II to allow for a shorter focal distance. Two Walimex Pro LED Strip Light Slim 300 Daylights were used for illumination. Laboratory imaging utilised a ZEISS Stereo Microscope Stemi 508 paired with a ZEISS Axiocam 208 colour digital camera for detailed morphological documentation.

### Sample preparations, imaging and analysis of histological sections

Specimens for morphological studies were euthanised under laboratory conditions using nitric acid solution (4.8 M HNO_3_, 300 mM NaCl in Milli-Q water) followed by fixation in 10 % formalin solution (4 % methanol stabilised-formaldehyde, 44.5 mM Na_2_HPO_4_ in Milli-Q water) for 2 days at room temperature (RT). The nitric acid solution formulation is the same as for the planarian fixative Steinmann’s fluid(54), except that it excludes mercury chloride (HgCl_2_) To prepare specimens, living individuals were placed in a 10 cm Petri dish containing a small volume of planarian water (PW)(55) or water from their habitat and allowed to stretch out. Once the specimens were fully stretched, nitric acid solution was poured over them and left for 30 seconds to 1 minute. After removing the liquid, the specimens were rinsed with 10 % formalin solution and transferred to 10 % formalin solution for two days at RT. Finally, they were transferred to 70 % ethanol in Milli-Q water using a soft brush or plastic pipette. The ethanol was replaced 2–3 times to ensure proper preservation. Samples were stored at RT. Toward the end of the project, we eliminated the formalin solution fixation from the protocol and samples euthanised with nitric acid solution were rinsed with 1x PBS and directly transferred to 70 % ethanol in Milli-Q water.

For histological sections, fixed specimens were dehydrated in a graded ethanol series, cleared in clove oil to facilitate the screening for the presence or absence of a copulatory apparatus, and embedded in Paraplast Plus in a Leica TP1020 automatic tissue processor. Serial sections 5 µm wide were made and stained with Mallory-Cason/Heidenhain stain(54). The stained sections were imaged with an Olympus VS200 widefield slide scanner with an Olympus UPLXAPO20X objective. Reconstructions of the copulatory apparatus were drawn manually on the basis of representative images using the Affinity Designer software and an XP-PEN Artist 15.6 Pro tablet. All processed slides will be deposited in the collections of the newly forming Biodiversitätsmuseum of Göttingen, Göttingen, Germany (ZMUG code).

### Metaphasic plate production and analysis

To prepare metaphase plates, individual *S. lugubris* specimens were cut twice between the root of the pharynx and the eyes. The middle piece was preserved in 100 % ethanol for barcoding analysis (see “Barcoding”). The posterior piece was fixed in nitric acid solution for histological studies (see “Sample preparation, imaging, and analysis of histological sections”). The anterior piece was used for karyological analysis. Anterior pieces were placed in 6-well plates containing PW(55) and incubated at 20°C. PW was replaced twice on the first day. After two days of regeneration at 20°C, samples were incubated for 6 hours in a solution of 400 ng/ml Nocodazole (Sigma, M1404) and 1 % DMSO in PW to elicit the metaphase-arrest of dividing cells. Specimens were then fixed in 3:1 methanol: glacial acetic acid and stored at −20°C for a minimum of 30 minutes, with some samples stored for several days before further processing.

Upon resuming the protocol, samples were brought to RT. Tissue containing the blastema and post-blastema regions was excised with a surgical razor blade, macerated in 45 % glacial acetic acid (diluted in Milli-Q water) for 30 minutes at RT, and the remaining tissue was discarded. The macerated tissue was minced on a glass slide, transferred to a 1.5 ml microcentrifuge tube, and pipetted up and down 10 times to create a cell suspension. A drop of this suspension was placed at the centre of a preheated (65°C on a hot plate) 15 mm Ø coverslip. After approximately 5 minutes of drying, dried coverslips were transferred to 12-well plates with the tissue side facing up and stored at 4°C.

To stain metaphase plates, coverslips were washed with 1x PBS for 5 minutes with gentle agitation, incubated for 30 minutes in a solution of 1.7 µg/ml of 4′,6-diamidino-2-phenylindole (DAPI) in 1x PBS (protected from light), and washed twice with 1x PBS and twice with Milli-Q water for 5 minutes each. Coverslips were gently dried with paper and mounted on glass slides using 10 µl of ProLong™ Glass Antifade Mountant (Thermo, Invitrogen, P36982). Mounted slides were cured for 12 hours at RT and stored at 4°C.

Metaphase plate images were captured using an Olympus VS200 widefield slide scanner equipped with X-Cite XYIS XT720L LED illumination. DAPI was excited using a 352–404 nm bandpass filter, and the fluorescence signal was collected with a 416–452 nm bandpass emission filter. Imaging involved three successive steps. A UPLFLN4X 4x air objective (NA = 0.13, WD = 17 mm) was used to locate the tissue area. Selected regions were scanned with a UPLXAPO20X 20x air objective (NA = 0.8, WD = 0.6 mm) to identify chromosome groups visually. High-resolution imaging of individual metaphase plates was performed with a UPLAPO100XOHR 100x oil objective (NA = 1.5, WD = 0.12), producing image stacks for each plate. Image stacks were processed in FIJI(56) and deconvolved using a theoretical point spread function (Gibson & Lanni 3D Optical Model)(57) for 200 iterations. A deconvolved substack of 1-3 planes was finally maximum projected to generate final metaphase plate images.

Chromosome counts were performed in FIJI. For Nottingham and Black Drim, we used laboratory populations derived from wild specimens maintained in our planarian collections(27). Four individuals and 4-7 metaphase plates per individual were analysed. For the Croatian population, five metaphase plates from a single individual were analysed. In total, 54 metaphase plates from 9 individuals were examined. Chromosomes were arranged in Figure 5 by relative length (%), calculated as (chromosome length*100)/total length of haploid complement.

### COI NCBI search

We retrieved all flatworm COI sequences from NCBI nucleotide archive present on 2024-04-11 using the following search term: “(((Platyhelminthes[Organism]) AND (CO1[Gene Name] OR COX1[Gene Name] OR COXI[Gene Name] OR COI[Gene Name] OR cytochrome oxidase subunit I[Title] NOT environmental[Title])))” and retrieved their taxonomic ID. The list of taxonomic IDs was further processed using TaxonKit (v0.18.0)(58) and plotted in R(59).

### COI degenerate primer design and barcoding

For designing primers for planarian COI, PlanMine transcriptomes(60, 61) and publicly available COI planarian sequences were aligned using Geneious(62). The species used for primer design belong to the families Dugesiidae (*Schmidtea mediterranea*, *Schmidtea polychroa*), Planariidae (*Planaria torva*, *Polycelis tenuis*, *Polycelis nigra*) and Dendrocoelidae (*Dendrocoelum lacteum).* The alignments were manually inspected for highly conserved regions suitable for primer design, with amplicons spanning roughly 880 bp of the planarian COI transcripts. Potential primer sequences were analysed using PerlPrimer(63) for potential hairpin formation and tuned in length for PCR annealing temperature optimisation. Primer positions with a degeneracy of three or fewer were included as equimolar wobble bases. To reduce the complexity of the primer mixture, higher degeneracy positions were replaced with the universal base inosine, which allows pairing with any base(64). Therefore, PCR with the MVCOI900 primer pair requires *Taq* polymerase for amplification, as proofreading polymerases, such as *Pfu,* cannot utilise inosine-containing primers(65). The MVCOI900 primers allowed direct Sanger sequencing of the PCR products (no need for cloning) (Additional File 1).

Genomic DNA was extracted from freshly cut pieces of living planarians or individuals fixed in 100 % ethanol with a combination of a guanidinium thiocyanate-based lysis buffer (GTC buffer) and phenol-chloroform extraction as previously described (66) (see “Rapid DNA isolation protocol”), with some modifications. Planarians were not demuconised. The lysis buffer was supplemented with 10 % v/v 1 M DTT right before use instead of 7 % v/v β-mercaptoethanol. Phase-maker gel tubes were replaced by 2 ml microcentrifuge tubes. Lysis and precipitation were done at RT. The pellet was washed twice with 1 ml 70 % EtOH to remove salts.

DNA sequences of the COI gene were obtained by PCR amplification for each individual. The final volume of each PCR reaction was 50 µl, including 2 µl of genomic DNA (1-20 ng), 5 µl of *Taq* Standard buffer, 2 µl of MgCl_2_ (25 mM), 1 µl of dNTPs (10 mM), 1.5 µl of each primer (10 µM), 0.25 µl of *Taq* polymerase (NEB) (Cat. No M0273S), and 36.75 µl of nuclease-free water.

The amplification conditions were the following: (1) 5 min at 95°C, (2) 30 s at 95°C, (3) 30 s at 50°C, (4) 1 min at 68°C, and (5) 3 min at 68°C. Steps 2, 3, and 4 were repeated for 30 cycles. Amplification products were visualised in a 1 % agarose gel and purified using QIAquick PCR Purification Kit (Cat. No 28104). The purified amplification products were sequenced in both directions using the amplification primers. Complementary sequencing reads were assembled using Geneious(62) and translated into amino acids to assure reading frame integrity. Alignments for barcoding were obtained using the software MAFFT (v7.490)(67) with default parameters implemented in Geneious(62).

### Haplotype networks

Two datasets were prepared to build haplotype networks and study the relationships among haplotypes of *S. lugubris*. Dataset 1 comprises an alignment of 16 COI sequences with a final length of 308 bp extracted from GenBank or generated for this study as described above. Dataset 2 includes 12 COI sequences with a final length of 702 bp exclusively generated in this study. In both cases, ambiguous positions (three in total) were removed from the original alignments before analysis. The aligned sequences were imported into DnaSP6 v.6(68) to generate Rohel Data files. Rohel Data files were imported into Network v10.2 to calculate haplotype networks with Median-joining network algorithm(69).

### RNA extraction, sequencing, and transcriptome assembly

RNA was extracted from planarian samples using a combination of TRIzol-based homogenisation, chloroform extraction, and commercial column purification. Briefly, for most samples, TRIzol reagent was added to the sample, which was then snap-frozen in liquid nitrogen and thawed on ice. The sample was homogenised using metal beads in the TissueLyser II, and chloroform was added to recover the upper phase. Isopropanol and high-salt-precipitation solution (0.8 M sodium-citrate and 1.2 M NaCl in Milli-Q water) were added to the aqueous phase, and the mixture was purified by the Direct-zol RNA MiniPrep (R2072) Kit.

A DNA-RNA combined extraction method was used for two samples (GOE00548 and GOE00576). Samples were homogenised in GTC buffer using the TissueLyser II, and the homogenate was separated into two phases using a phenol-chloroform method. The upper phase was transferred to two new tubes, with 40 % of the volume used for DNA extraction and 60 % for RNA extraction. DNA samples were treated with *RNase A* and purified using a standard DNA extraction protocol (see above). RNA samples were purified as described above.

RNA quality was assessed using a Bioanalyzer RNA 6000 Nano kit (5067-1511). Double-indexing was used to minimise cross-contamination of transcriptomes. RNA was then processed for 150 base pair paired-end Illumina sequencing, performed by the Dresden Concept Genome Center and Azenta Life Sciences.

Adapters, low complexity reads, and low-quality bases were removed from the short reads using fastp (v=0.23.4, ‘-l -c -5 5 -M 30 -r 5 -l 35 --detect_adapter_for_pe’)(70). Then, Kraken 2 (v=2.1.3)(71) was used with the standard database and default parameters in combination with KrakenTools(72) (‘extract_kraken_reads.py –exclude –taxid 2 2157 10239 –include-children’) to filter out potential contamination from viruses, bacteria, and archaea. De novo transcriptome assemblies were generated from the filtered reads using Trinity (v=2.15.2, ‘--SS_lib_type RF‘)(73).

Potentially contaminating contigs originating from the bovine liver food or the experimenters themselves were removed using the mmseqs taxonomy tool ‘--taxon-list ‘33208&&!9913&&!9606’. Protein coding sequences were predicted using TransDecoder (v5.7.1, ‘-m 75 –single_best_only’) (https://github.com/TransDecoder/TransDecoder), requiring a minimum length cut-off of 75 amino acids. Open reading frames (ORF) below the cut-off were included in case of significant homology scores identified using an MMseqs2 profile search against Pfam, eggNOG, or BLAST search against Swiss-Prot.

Finally, the contig with the longest, complete protein prediction from each De Bruijn graph component was chosen for subsequent analysis. In cases where only incomplete protein predictions were obtained, the contig with the longest predicted protein was chosen.

### Phylotranscriptomics

We supplemented the five transcriptomes generated in this study with transcriptomes extracted from(27, 74) and performed gene supermatrix construction as described in(27). Briefly, for each transcriptome, the longest protein per transcript was predicted using TransDecoder (v5.7.1, ‘-m 75 – single_best_only’), homologous groups were inferred using OrthoFinder(75) (v2.5.4, -M ‘msa’ -I 1.5 -z -ot -X), a multiple sequence alignment for each homologous group was created using MAFFT (v7.525)(67), and a gene tree was inferred using FastTree (v2.1.10)(76). To arrive at a set of orthologous sequences for phylogenetic inference, the homologous groups were pruned using PhyloPyPruner (https://pypi.org/project/phylopypruner/) with midpoint rooting and ‘trim_lb 4, min_taxa=25, Maximum inclusion, min_otu_occupancy = 0, min_gene_occupancy = 0’. The final supermatrix consisted of 1701 protein alignments from 55 transcriptomes with 1069932 columns, 784087 distinct patterns 552857 parsimony-informative, 180422 singleton sites, 336653 constant sites and an overall percentage of missing data of 55.9 %. The species tree was then inferred using IQ-TREE(77), first determining the best substitution model for each partition using ModelFinder (‘-m TEST’) and calculating 1000 ultra-fast bootstraps to assess branch support (-bnni -bb 1000). Trees for COI were inferred with IQ-TREE(77) with the same settings as described above.

## Results

### DNA barcoding analysis reveals taxon biases in planarian COI sequences available in GenBank

Toward our goal of facilitating the species-level identification of planarians, we first assessed the taxon coverage of COI reference sequences currently available in GenBank (National Center for Biotechnology Information, NCBI)(78). Specifically, we compared the number of GenBank COI accessions for each major group within the phylum Platyhelminthes to the number of known species per group recorded in the Turbellarian database (see Methods for details)(79). Out of 41743 platyhelminth COI entries, 87.6 % were of parasitic Neodermata, which include medically relevant species like *Schistosoma mansoni* and, therefore, likely reflect a positive sampling bias (Fig. 1A). The remaining entries were split between Catenulida (0.1 %) and the free-living Rhabditophora (excluding the parasitic Neodermata) (12.2 %). Among those, Tricladida (planarians) comprised 74.7 % of COI entries, a disproportionately high representation compared to other groups such as Polycladida (10 % of entries) considering the number of species (Fig. 1A). Within Tricladida, 55.5 % of accessions belonged to Geoplanidae (land planarians, 55.4 % of the known Triclad species). In comparison, freshwater Continenticola constituted 44.5 % of the total number of accessions: 37.8 % for Dugesiidae (13.1 % of the known triclad species), 6 % for Planariidae (10.3 % of the known species), and only 0.6 % for Dendrocoelidae (14.5 % of the known species) (Fig. 1A). Notably, no COI sequences were available for Kenkiidae, a freshwater family with 24 described species, and only a single accession representing Cavernicola (nine species) was present in the data set. Finally, Maricola, a primarily marine triclad group with 83 known species, was represented by only eight accessions. Thus, the number of COI accessions is disproportionately high for the family Dugesiidae and disproportionately low for other freshwater groups like Planariidae or, particularly, Dendrocoelidae.

**Figure 1.**
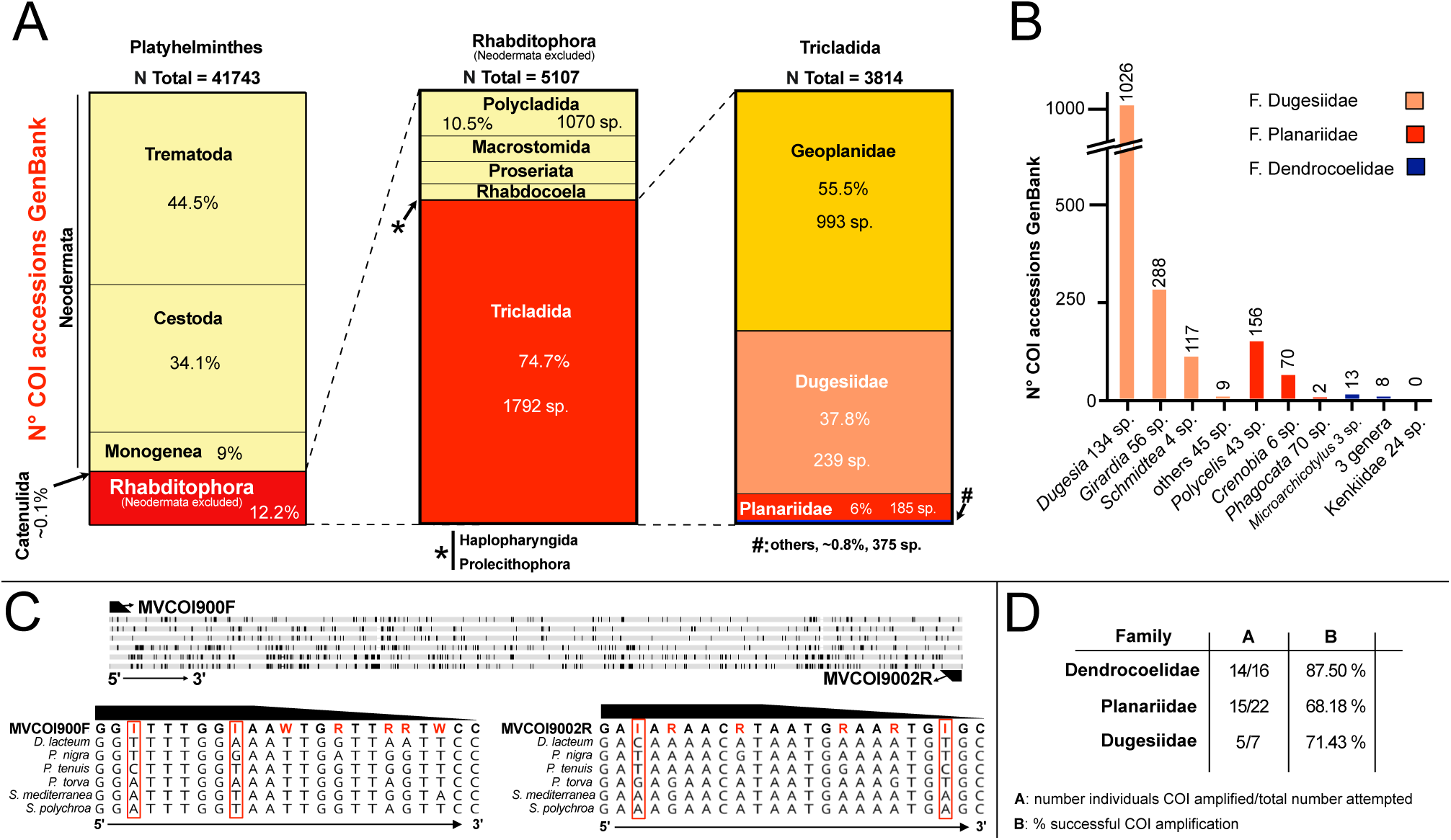
Analysis of the GenBank COI sequence accessions library indicates an underrepresentation of specific planarian groups. **A**. GenBank COI accession in three taxonomic groups: Platyhelmintes (left), Rhabditophora excluding Neodermata (middle), and Tricladida families (right). The total number of accessions (N Total), the percentage they represent from the total per group (%), and the approximate number of species (sp.) are indicated for relevant groups and subgroups. **B.** Number of GenBank COI accession for selected genera and approximate number of species (sp.). **C.** Degenerated primer design. Top, design of the new COI primer pair MVCOI900 on basis of an alignment of the COI gene sequences from six planarian species. Primer binding positions are indicated. Black bars: variable sequence positions. Bottom, sequence alignment detail in the primer binding site. Primer highlighted in bold. Red: variable base positions in the primer. Red squares: highly variable positions represented by an inosine (I) base in the primer. For MVCOI900_F, the first “W” wobble base in the primer sequence stems from an initially less stringent selection of sequences during primer design. **D.** Primer COI gene sequence amplification success by PCR.

The genus *Dugesia* comprises 56.1 % (134 sp.) of the known Dugesiidae species but accounts for 71.3 % of the family accessions, indicating a notable overrepresentation. *Girardia*, which comprises 23.4 % (56 sp.) of Dugesiidae species, constitutes 20 % of accessions, while *Schmidtea*, with only four species, contributes disproportionately with 8.1 % of accessions (Fig. 1B). In contrast, the remaining nine genera, collectively encompassing 13.8 % of species, account for only 0.6 % of accessions (Fig. 1B). These findings reveal significant biases favouring *Dugesia* and *Schmidtea* within the Dugesiidae family. This trend of overrepresentation is mirrored in other planarian families. For example, within Planariidae, *Polycelis* and *Crenobia* are disproportionately well-represented, while several other genera are sparsely included in GenBank. These patterns within the currently available COI reference sequences suggest systemic biases in taxonomic sampling, which may become even more apparent with finer-scale analyses at the species level.

Overall, our results confirm the existence of pronounced taxonomic biases at both family and genus levels in current planarian barcode records in GenBank, which makes barcode-based species identification challenging. These findings highlight an urgent need for systematic efforts to integrate COI barcode data from undersampled taxonomic groups into genetic repositories.

### Optimisation of COI primer pair for European freshwater planarians

One reason for the observed coverage biases in the COI GenBank record is that the available primer pairs perform poorly on planariids and dendrocoelids (M. Riutort, M. Vila-Farré; unpublished observations). Towards the goal of improving primer performance and COI taxon coverage, particularly for European species of planarians, we used the COI gene sequence representation in our collection of *de novo* assembled planarian transcriptomes(27). Sequence alignments of representative species belonging to three key European freshwater families (Dugesiidae, Planariidae and Dendrocoelidae) highlighted significant polymorphisms within the primer binding region. To accommodate these SNPs, we designed a new primer pair (MVCOI900) that includes degenerate base positions or inosine as a universal base at highly polymorphic positions(80, 81) (Fig. 1C). MVCOI900 amplifies ∼880 bp in the coding region of the COI gene (Fig. 1C; Additional File 1). We tested the primers’ amplification efficiency on phenol-chloroform DNA preparations from a range of planarian species, as commercial DNA isolation kits perform poorly on planarians (see Methods). Amplification success rates with our standardised protocol were 87.5 % for dendrocoelids, 71.4 % for dugesiids, and 68.18 % for planariids (Fig. 1D). Within planariids, amplification was challenging for *Phagocata* or *Crenobia*, possibly indicating species- or genus-specific variability in primer efficiency and/or species-specific PCR inhibitors(82). Even though generating a truly universal planarian barcoding primer will require further optimisation, the MVCOI900 primer pair allows a significant taxon expansion and, thus, a means of addressing the previously noted COI coverage bias across planarian taxa.

### Croatia field sampling campaign and preliminary classification of the planarian samples

To evaluate the utility of the new primers on field-collected material and to explore the diversity of the planarian fauna of Croatia, we sampled freshwater habitats in the country’s karst region, including rivers, lakes, and springs (for sampling methodology, see(27) and methods section) (Fig. 2aA). The collected planarians were coarse-classified at the genus or species level based on external morphological features of live specimens (Fig. 2aB) guided by our taxonomic expertise and the existing literature on European and Western Balkan planarians(43–51, 83, 84).

**Figure 2a.**
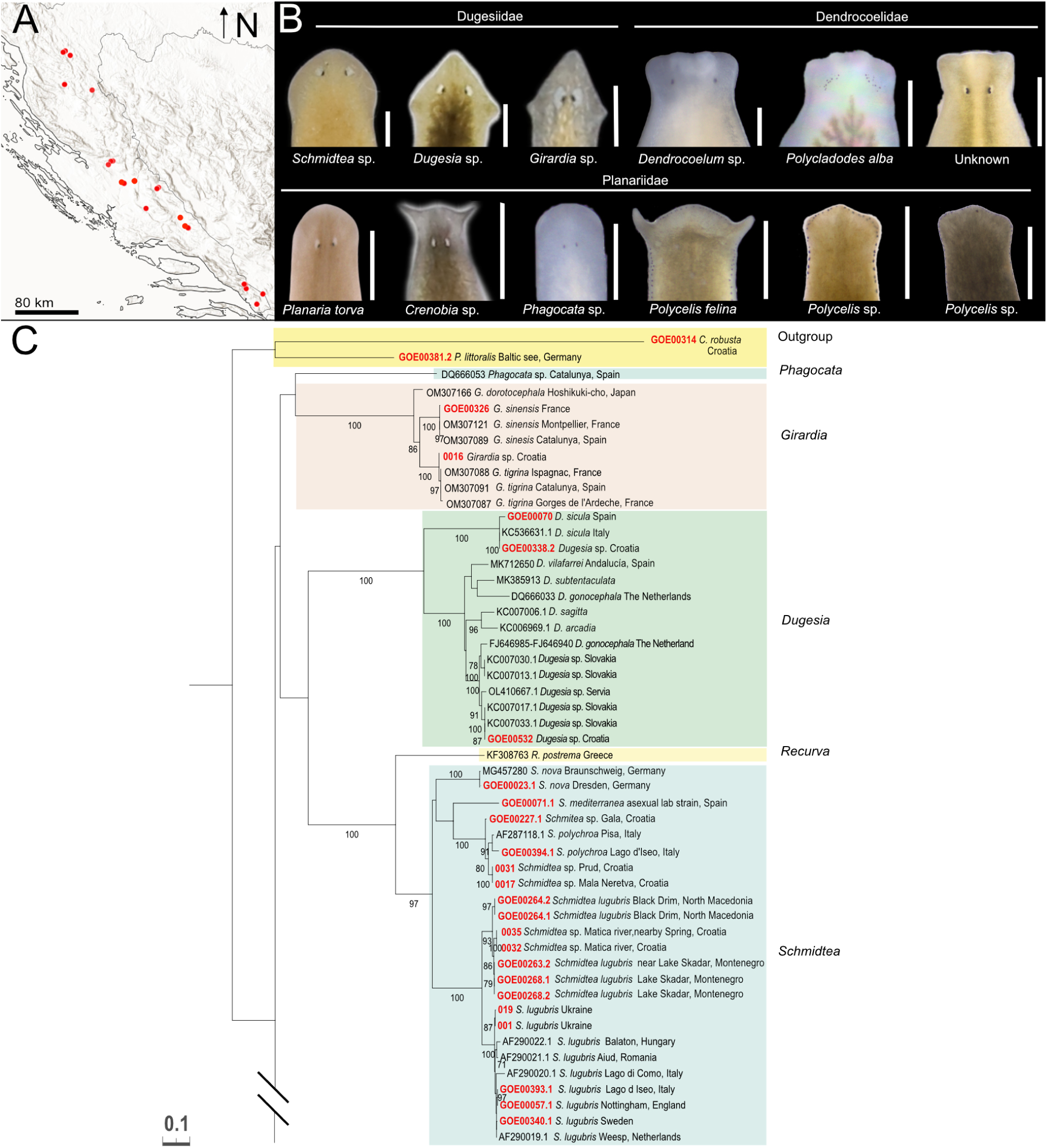
Barcoding of the Croatian planarians, part 1. **A.** Field sampling sites with planarian occurrences. The sampling site at Ombla spring, north-east of Dubrovnik, is not indicated on the map. **B.** Preliminary taxonomic assignment of the collected specimens, based on morphological traits of head anatomy. Scale bar, 1 mm. **C.** First part of the tree obtained with IQ-TREE from “Dataset Barcoding” containing the indicated outgroups, dugesiids and one *Phagocata* sample. The tree continues in Figure 2b. The numbers at the nodes represent the bootstrap values. Red terminals correspond to individuals sequenced in this study. Colour boxes delimit planarian genera.

The morphological characteristics used for genus identification included, among others, the number and arrangement of eyes relative to the body margin and head shape (Fig. 2aB illustrates the variability of this character), dorsal and ventral body colouration, number of visible pharynges and shape and type of movement of the body margin. Our initial classification identified *Schmidtea* sp. (five localities), *Dugesia* sp. (two localities), and *Girardia* sp. (one locality) among dugesiids; *Dendrocoelum* sp. (11 localities), *Polycladodes alba* (one locality), and an unidentified pigmented dendrocoelid (three localities); and the planariids *Phagocata* sp. (three localities), *Planaria torva* (three localities), *Crenobia alpina* (one locality), *Crenobia montenigrina* (one locality), *Polycelis felina* (eight localities), and *Polycelis* sp., with auricles resembling those of *P. nigra* and *P. tenuis* (eight localities) (Fig. 2aB).

The unidentified pigmented dendrocoelid mentioned above was particularly interesting, as the only known pigmented dendrocoelid species in Europe outside the Ohrid and Prespa lakes region is *Bdellocephala punctata*, which possesses a more distinct adhesive organ and different dorsal pigmentation. The Ohrid and Prespa lakes region is home to various pigmented *Dendrocoelum* species(85). The collection of these remarkable specimens, approximately 600 km north of the Ohrid-Prespa region, highlights the scientific interest of Croatian triclads.

### DNA barcoding of Croatian planarians

To further characterise the collected specimens, we generated COI barcodes using our optimised primers for the Croatian samples and others collected during other European lab expeditions. Furthermore, we included existing GenBank COI sequences of European freshwater planarians. The final dataset (“Dataset Barcoding”) contains 124 sequences, including 77 new sequences generated during the course of this study via amplification with the MVCOI900 primers (except for two generated with a different primer pair; Additional File 2). The length of the amplicons ranged from 734 to 890 bp, with a mean of 844 bp (S.D.= 36.6), and the 47 existing GenBank records of European planarians varied in length from 302 to 1755 bp, with a mean of 620 bp (SD= 286.2). Notably, the primers also amplified two maricola, *Procerodes littoralis* and Croatian individuals of *Camerata robusta*, which is only the second known population of this species(86), that we used as outgroups in the subsequent analyses. The significant expansion of planarian COI barcoding sequences in GenBank again highlights the utility of the new MVCOI900 primer set. All new sequences will be deposited in GenBank, with accession numbers provided in Additional File 2.

To infer the species identity of our samples, we generated a phylogenetic tree from “Dataset Barcoding” (Fig. 2aC, 2b). We compared the similarity of our unidentified samples with the identified samples in the dataset. Our tree includes poorly investigated planarian genera, e.g., *Dendrocoelum*, *Polycladodes*, European *Polycelis* or *Planaria*, thus representing a step forward in planarian barcoding. Analysis of the barcoding tree reveals that the traditionally recognised genera are recovered as monophyletic, frequently with very high statistic support, with two exceptions from GenBank sequences that are placed in positions inconsistent with their genus assignation: *Phagocata* sp. (GenBank: DQ666053) clusters with *Girardia* while *Dendrocoelum lacteum* (GenBank: AF178312) clusters with *Bdellocephala punctata*. In the original publication(87), *Dendrocoelum lacteum* (AF178312) clusters with *Polycelis*, but the COI tree does not include *B. punctacta* or other *Dendrocoelum* species (Fig. 6.5), a situation later repeated in(88) (see Supplementary data 2D). Interestingly, in the original publication(88), *Phagocata* (DQ666053) is placed next to other planariids in a position coherent with the genus designation, far from *Girardia* (see Supplementary data 2D). Given that other *Phagocata* and *Dendrocoelum* sequences cluster cohesively with different samples of their genus in our tree, the taxonomic assignment of DQ666053 and AF178312 in GenBank is very likely erroneous. Further investigation is required, particularly considering the extreme scarcity of *D. lacteum* COI accessions in GenBank. We selected sequences with GenBankID DQ666033 (generated in (88)) and FJ646985-FJ646940 (see Additional file 2) to represent the species *Dugesia gonocephala* in our COI dataset. However, these sequences cluster independently within the *Dugesia* clade in the tree, suggesting a problem with species designation.

**Figure 2b.**
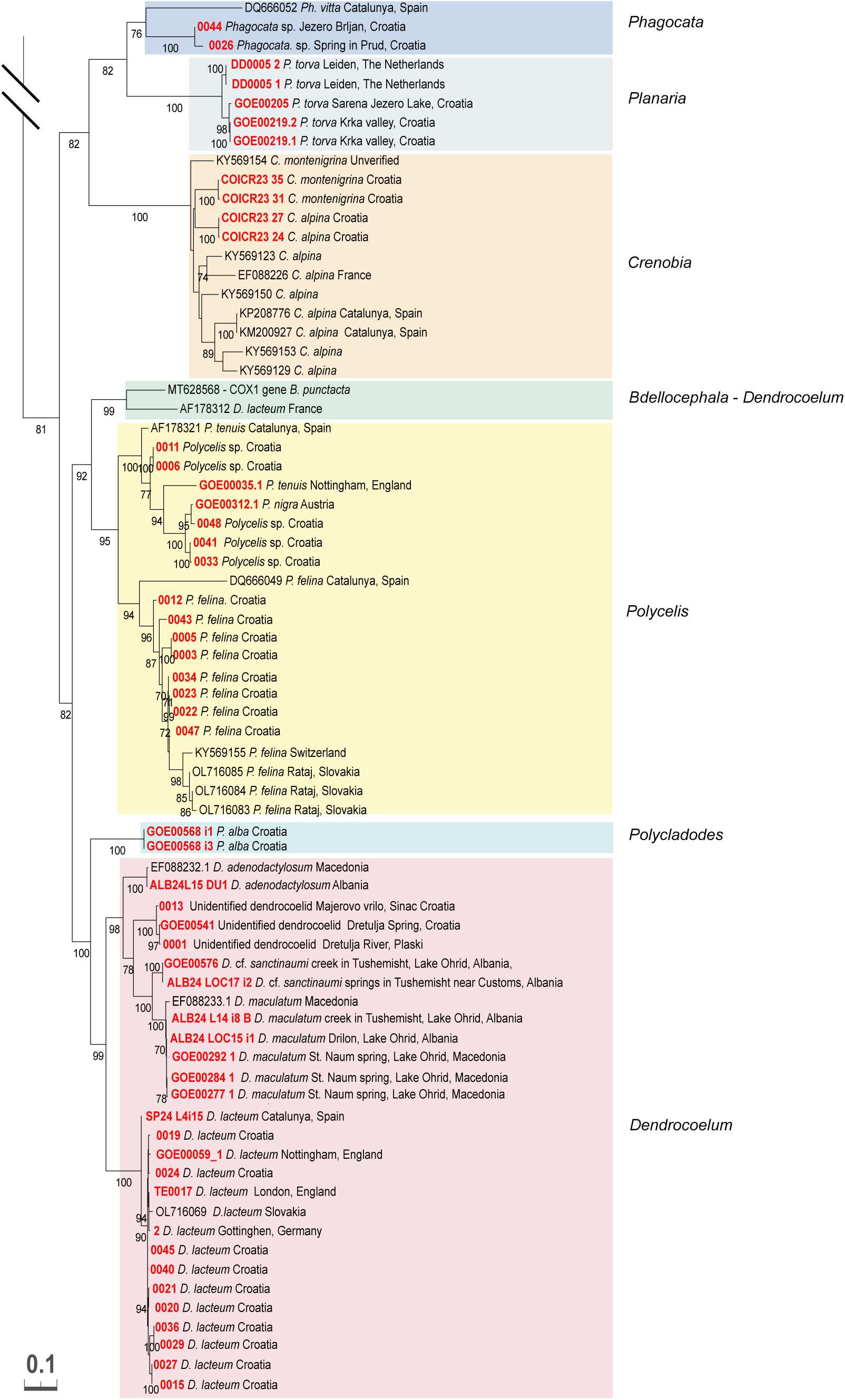
Barcoding of the Croatian planarians, part 2. Second part of the tree obtained with IQ-TREE from “Dataset Barcoding” containing the planariids and dendrocoelids (See Figure 2a).

Several Croatian samples were identified through matches in GenBank or our dataset. Among the dugesiids collected in Croatia, we positively identified *Girardia tigrina*, *Dugesia sicula*, *Schmidtea polychroa* and *Schmidtea lugubris* (Fig. 2aC, 3). We also successfully identified the dendrocoelid *Dendrocoelum lacteum* and the planariids *Planaria torva* and *Polycelis felina* (Fig. 2b). Intriguingly, the tree topology revealed two sister clades of *Schmidtea lugubris*: one comprising samples from various European localities and another containing exclusively samples from the Adriatic area (from Croatia in the North to the Ohrid lake in the South) (Fig. 2aC). This divergence warrants further investigation to clarify the taxonomic status of the two clades (see below).

**Figure 3.**
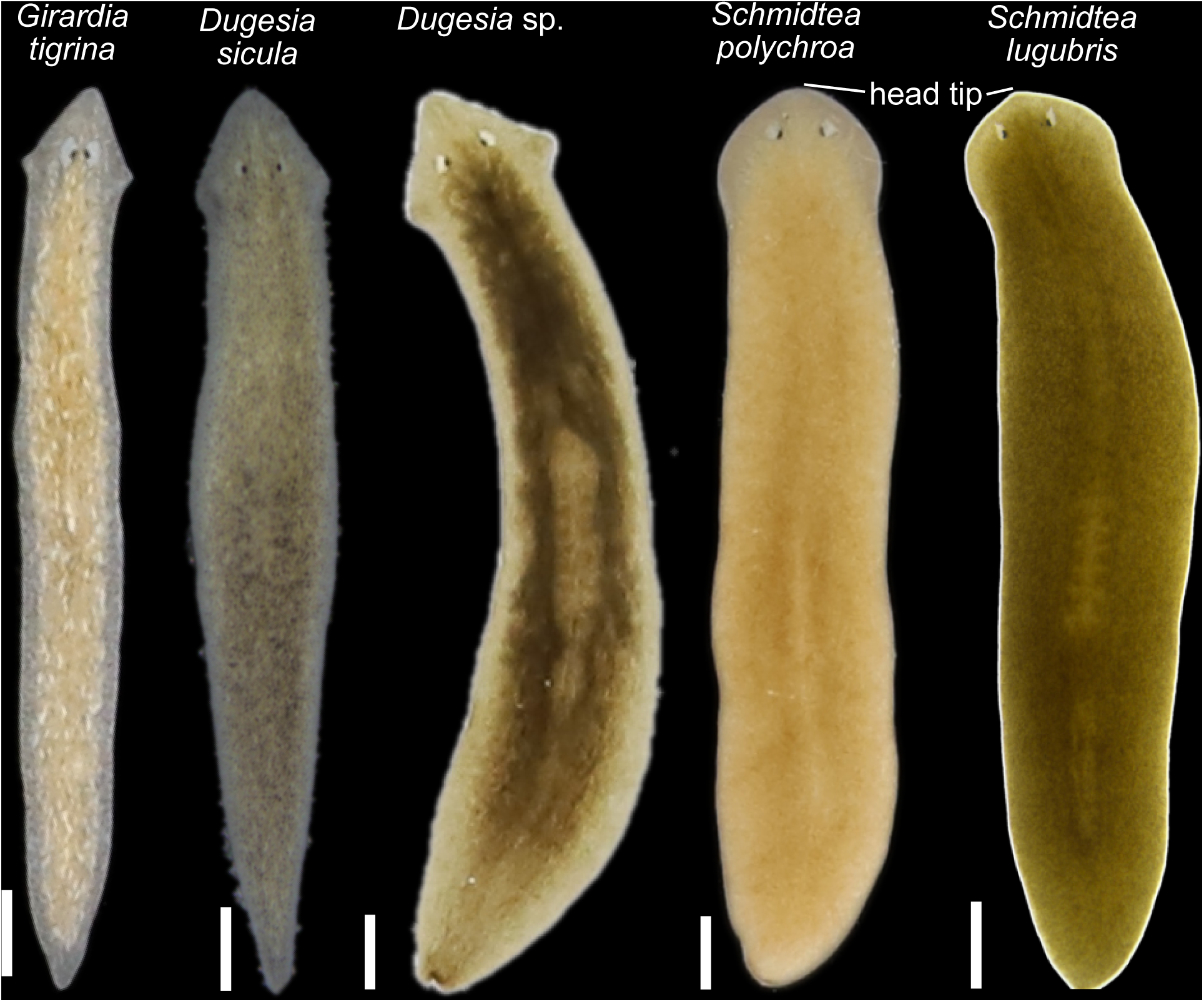
Diversity of dugesiids. **A.** Living planarian specimens representing the dugesiids collected during our expeditions. Scale bar, 1 mm.

Barcoding also provided clues for unresolved samples. For instance, GOE00532 (*Dugesia* sp.) (Fig. 2aC) is likely *D. gonocephala* or a related form. However, due to the lack of identified close matches, we considered it advisable to classify individuals from the same population using classical taxonomy.

Individuals assigned to *C. montenigrina* and *C. alpina* (corresponding to two individuals from a single population per species) clustered in two sister clades, each corresponding to each species, nested in a clade with other *Crenobia* (Fig. 2b). Since the single GenBank accession for *C. montenigrina* clustered separately from the Croatian *C. montenigrina*, the specimens were further investigated with classical taxonomic methods to clarify their species’ assignment (see below).

Barcoding was inconclusive for the two *Polycladodes alba* accessions, two *Phagocata* sp., several *Polycelis* sp. samples, and three pigmented dendrocoelids (Fig. 2b). The *Polycladodes alba* and Croatian *Phagocata* sequences had no closely related matches in GenBank. Meanwhile, the three pigmented dendrocoelids clustered with members of the genus *Dendrocoelum*, including pigmented species from Lake Ohrid, suggesting they likely belong to this genus. Interestingly, the three individuals of pigmented dendrocoelids (each one from a different locality) are grouped by geographic proximity into two clusters (Fig. 2b).

Overall, this analysis again highlights the fragmentary record of COI accessions in GenBank for many widespread European species (e.g., *P. torva*, three *Polycelis* species, *D. lacteum*, or *P. alba*) and the difficulties that result from barcoding-based species identification.

### Phylogenetic relationships of the Croatian dendrocoelids and transcriptomic comparison of the *Schmidtea lugubris* clades

To obtain further insights into the phylogeny of Croatian dendrocoelids and the two *Schmidtea lugubris* clades revealed by our barcoding effort, we generated de-novo assembled transcriptomes via our pipeline(60, 61) of *Polycladodes alba*, the pigmented dendrocoelid, *Dendrocoelum* cf. *sanctinaumi* (a pigmented form from the Ohrid region), *Dugesia* sp. (that we plan to include in future studies), and a Croatian population of *S. lugubris*. Together with the recently published transcriptomes of four *Schmidtea* species(74) and the transcriptomes published in(27), this amounted to a dataset of 55 transcriptomes (51 planarians and four outgroups). Of note is that two transcriptomes in the tree also correspond to Croatian population of *S. polychroa* and *P. torva,* which we published previously(27, 74). We extracted broadly conserved orthologues(73, 75, 77) and constructed a phylogenetic tree (see Methods). The resulting phylogeny achieved maximum branch support for all but one node (Fig. 4). Consistency between our phylogeny and that in (27) (species common to both trees appear in a similar position forming similar phylogenetic clades) underscores the robustness of the tree topology and validates the inclusion of new transcriptomic data.

**Figure 4.**
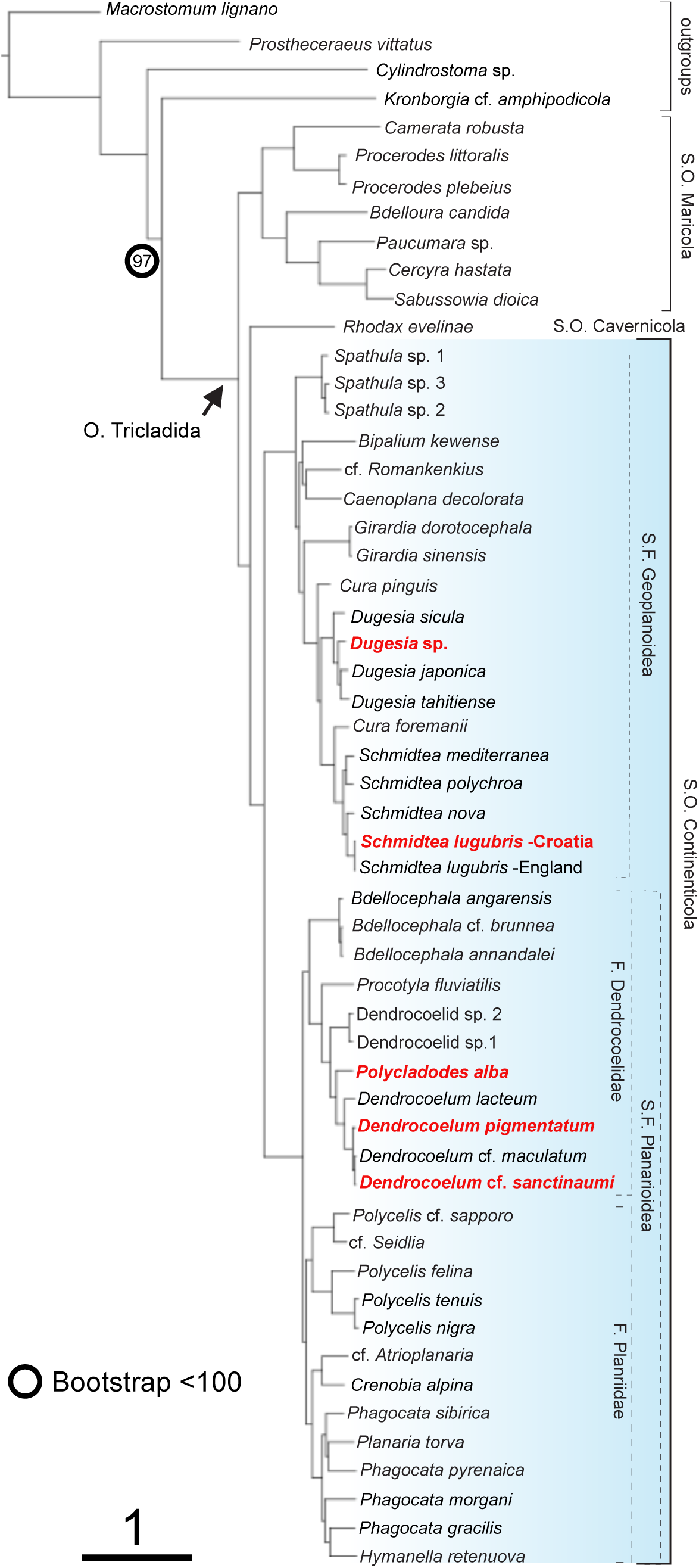
Phylogenetic relations of Croatian dendrocoelids and *S. lugubris*. Transcriptome-based IQ-TREE with transcriptomes de-novo generated for this paper highlighted in red. Colour shading marks the Continenticola, which contains the freshwater groups analysed. Bootstrap values 100, except where indicated.

The taxonomic status of *Polycladodes* has been debated, with some earlier publications considering it a subgenus of *Dendrocoelum*(47). We adopt the use of the genus name *Polycladodes* Steinmann, 1910, over *Dendrocoelum* Örsted, 1844, for *P. alba*, following recent taxonomic classifications(89, 90). Our analysis positions *Polycladodes alba* as a sister group to the four *Dendrocoelum* species included in the phylogeny (*D. lacteum*, two pigmented species from Lake Ohrid) and the pigmented dendrocoelid from Croatia(Fig. 4). As expected, this topology supports a close evolutionary relationship between *Polycladodes* and *Dendrocoelum*, while also affirming the genus-level distinction of *Polycladodes*.

In our phylogenetic tree, the Croatian pigmented dendrocoelid forms a clade with two other pigmented species from Ohrid, supported by maximum branch support and with relatively shallow branch lengths, indicating a close relation among the species (Fig. 4). This result suggests that it is a member of the genus *Dendrocoelum*, in agreement with the barcoding data, and highlights the likely shared evolutionary history of pigmented *Dendrocoelum* species in the region that will be elaborated in the general discussion.

To understand whether the two clades of *S. lugubris* (Fig. 2aC) are different enough to constitute separate species, we compared the branch length distance between the sister species of *Schmidtea* and the two *S. lugubris* forms. Our tree confirms the established relationships within *Schmidtea*(74, 91) (Fig. 4): *S. mediterranea* is sister to *S. polychroa,* and *S. lugubris* is sister to *S. nova*. The branch length distances between *S. mediterranea* and *S. polychroa* (0.1293) and between *S. nova* and *S. lugubris* (British population) (0.1359) are comparable. In contrast, the distance between the two *S. lugubris* clades (0.0121) is approximately ten times smaller, suggesting significantly lower divergence. This lower divergence in the phylogenetic tree, together with the differentiation into clades in the COI barcoding tree (Fig. 2aC), points to the existence of previously unknown intraspecific variability and genetic structure rather than the split into two species of *S. lugubris*.

We constructed two COI datasets with sequences of different lengths to investigate the genetic diversity of the two *S. lugubris* clades, Dataset 1 (308 bp) and 2 (702 bp) (see Methods). We identified 10 haplotypes in Dataset 1 and 8 in Dataset 2. The haplotype networks separate two well-differentiated groups (Fig. 5A, Additional File 3) that we named the Central and Adriatic clades. In Dataset 1, the Central clade encompasses individuals distributed across central Europe, with minimal divergence across a large area (from the Netherlands to Sweden, the United Kingdom, and northern Italy), as well as populations in the east regions of Europe forming a more complex network (Italy, Hungary, Romania, and Ukraine) (Fig. 5A, B). Individuals in the east of the Central clade exhibit no shared haplotypes with the Western individuals or among them. The Adriatic clade consists of three haplotypes in Dataset 1, restricted to Croatia, Montenegro, and North Macedonia. The Adriatic clade displays minor genetic differentiation between the sampled individuals in this dataset.

**Figure 5.**
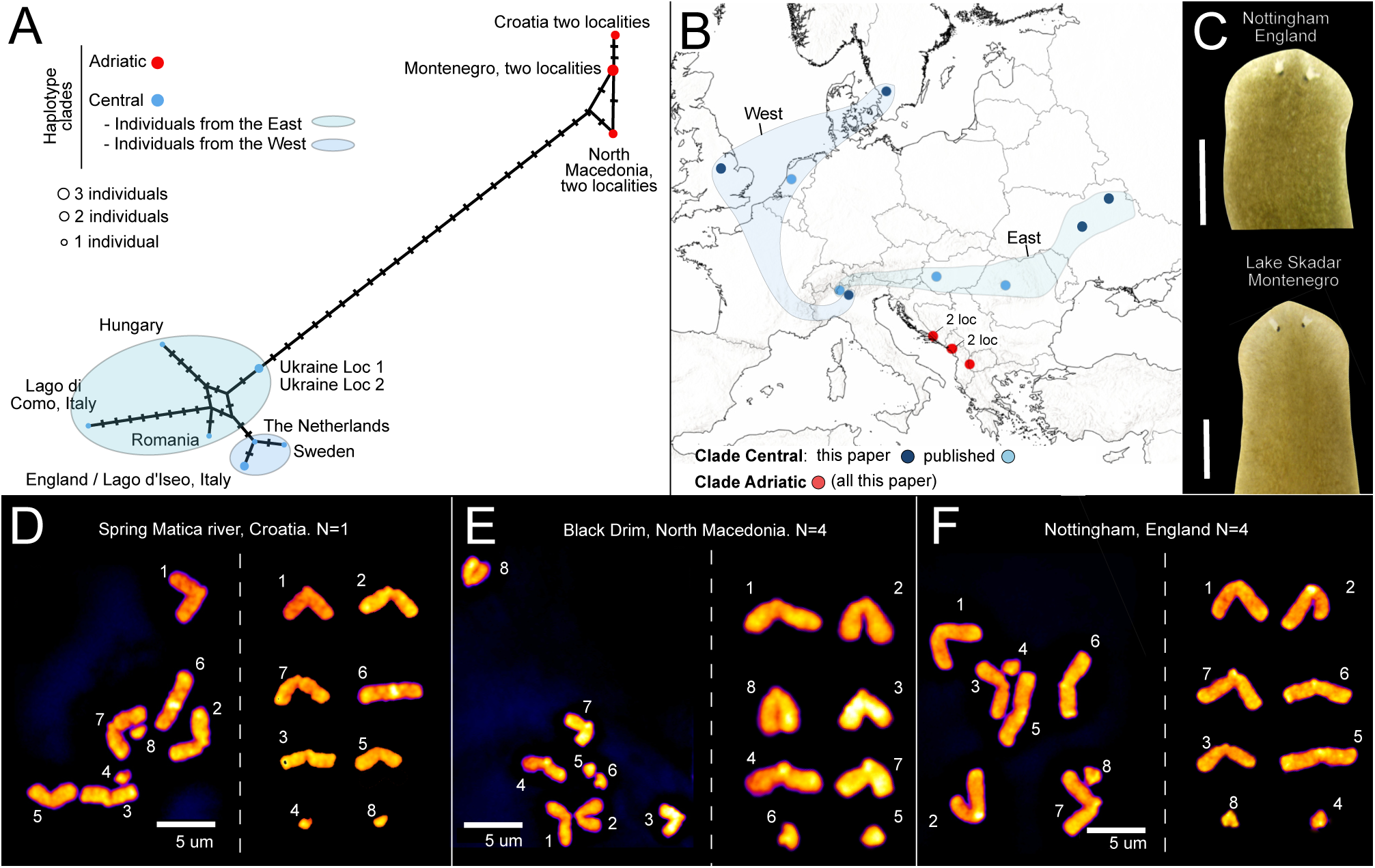
A. The Adriatic populations of *S. lugubris* form a distinct genetic clade. **A.** Haplotype network for the mitochondrial gene COI from Dataset 1 (sequence length 308 bp). Each circle represents a different haplotype, and the size of the circle indicates the frequency of each haplotype. Black bars represent intermediate (non-present) haplotypes, and lines connecting haplotypes (existing or not) represent one nucleotide change. The colour scheme is used to differentiate the major haplotype clades (see legend) and the origin of the individuals sampled within the central clade. **B.** Distribution map of the two haplotype clades. Each dot represents a locality. Dots representing two nearby localities are indicated (2 loc). Curves enveloping localities correspond to the Eastern and Western localities described in **A** for the central clade. The origin of COI sequences used to assign each locality to a clade (published data or this study) is indicated. **C.** Living specimens showing the characteristic pointed head of *Schmidtea lugubris.* **D-F.** Metaphasic plate (left) and chromosome complements arranged in pairs (right) of individuals from three populations arranged in pairs. Numbers indicate the correspondence between chromosomes. Individuals from the three populations show diploid karyotypes (2n = 8) with three acrocentric and one small chromosome. The number of individuals analysed per population (N) is indicated next to the population name.

Conversely, in Dataset 2, the Adriatic clade gains haplotype diversity (from three to four haplotypes), and the three areas of origin are better differentiated, indicating the existence of interesting diversity in the Adriatic zone as well. Long sequences from the Central clade are scarce, resulting in only four haplotypes in Dataset 2. Notably, the Ukrainian individuals remain distinct from the Western individuals. In summary, our analysis consistently revealed the existence of two highly distinct genetic clades within *S. lugubris*.

### Systematic section

To address some of the ambiguities in our barcoding analysis, we conducted a detailed histological analysis of individuals from populations where enough material was available, a standard method in triclad taxonomy(33, 51, 85). *Planaria torva* was included to curate the submitted COI accession as the first GenBank COI accession of this species. The systematic section as a whole presents data for five taxons in three families: the dugesiid *Schmidtea lugubris*, the dendrocoelids *Polycladodes alba* and the pigmented dendrocoelid (described as a new species, *Dendrocoelum pigmentatum*), and the planariids *Planaria torva* and *Crenobia montenigrina*.

**Order: Tricladida Lang, 1884**

**Suborder: Continenticola Carranza, Littlewood, Clough, Ruiz-Trillo, Baguñà & Riutort, 1998**

**Family: Dugesiidae Ball, 1974**

**Genus: *Schmidtea* Ball, 1974**

**Species: *Schmidtea lugubris* (Schmidt, 1861)**

**Material Examined**. H0273, Matica river, nearby Spring, Dusina, Croatia, (43.1762° N, 17.4069° E), 15/09/21, coll. Jochen C. Rink, Miquel Vila-Farré, Ludwik Gąsiorowski, Uri Weill and Rick Kluiver, sagittal sections on 20 slides. H0274, ibid., horizontal sections on 10 slides. H0275, ibid., sagittal sections on 10 slides. H0802, laboratory population originally collected from Nottingham, United Kingdom (52.942432° N, −1.113739° E), 2011-12, coll. Jochen C. Rink, sagittal sections on 12 slides. H0803, ibid., sagittal sections on 8 slides. H0804, ibid., sagittal sections on 7 slides. H0805, laboratory population originally collected from Lago d’Iseo, Italy (45.724999° N, 10.053001° E), 2013, coll. Miquel Vila-Farré, sagittal sections on 9 slides H0806, ibid., sagittal sections on 7 slides H0807, ibid., sagittal sections on 7 slides. H0813, laboratory population originally collected from Lake Skadar, Karuč, Montenegro (42.35791° N, 19.10661° E), 2014, col. Jochen C. Rink, Miquel Vila-Farré, Helena Bilandžija, sagittal sections on 11 slides. H0814, ibid., sagittal sections on 11 slides. H0815, ibid., sagittal sections on 9 slides. H0816, laboratory population originally collected from Crni Drim (Black Drim, Ohrid outflow), Republic of North Macedonia (41.355435° E, 20.623436° N), 2014, col. Jochen C. Rink, Miquel Vila-Farré, Helena Bilandžija, sagittal sections on 23 slides. H0817, ibid., sagittal sections on 15 slides. H0818, ibid., sagittal sections on 11 slides.

### Taxonomic discussion

Wild Croatian *S. lugubris* individuals measure up to ∼1.2 cm in length and present a uniformly brown dorsal pigmentation. Other strains of *S. lugubris* maintained in our collection(27) are mostly brown. The characteristic but challenging-to-detect pointed head of *S. lugubris*(92), which contrasts with the rounded heads of *S. polychroa*, is visible in some individuals from both the Central and the Adriatic clade (Fig. 3, 5C). The copulatory apparatus anatomy conforms with that expected for *S. lugubris*. They present two seminal vesicles, the first narrow in the middle and broader in its upper and lower sections and coated with a well-developed layer of circular and longitudinal muscles. The narrow ejaculatory duct emerges from an area close to the ventral section of the seminal vesicle. It later widens to form the second vesicle, which is smaller, providing a strong muscle layer that combines longitudinal and circular muscles. Once in the very large penis papilla, the ejaculatory duct runs initially centrally to bend markedly in the middle section of the papilla to run again centrally to open into a very long and distinctive nipple characteristic of the species (Additional File 4). Of note, several individuals from the United Kingdom and one from North Macedonia (H0802, H0803, H0804 and H0818) present an expansion in one of their vasa deferentia. The copulatory apparatus of *S. lugubris* in individuals of other localities is similar: the penis bulb and the two seminal vesicles are surrounded by very muscular tissue, the ejaculatory duct follows a similar trajectory, the penis papilla is large and presents a nipple that, although variable in length, is presented in all the individuals analysed (Additional File 4, 5).

### Karyological analysis

Karyological studies clarified the complex taxonomic history of the four *Schmidtea* species by identifying seven chromosomal strains (biotypes) (see(91) for a summary). *S. lugubris* corresponds to biotype E, diploid with a karyotype (2n = 8) and three acrocentric (terminal or nearly terminal centromeres) and one submetacentric chromosome(91, 93). Our analysis revealed similar karyotypes in Adriatic and Central clade individual. All metaphase plates analysed (a total of 54; see methods) showed eight chromosomes except for five plates, which exhibited 7, 9, or 12 chromosomes. Among the eight chromosomes, six were acrocentric while the small size of the remaining two chromosomes precluded counting their arms. This karyotype 2n = 8, with a haploid complement of three large acrocentric and one small chromosome (Fig. 5D-F) aligns with biotype E, confirming the specimens as *S. lugubris*. Thus, our analysis, integrating a multigene phylogeny, barcoding, morphological, and karyological data, strongly suggests that the Adriatic and Central clades correspond to distinct genetic lineages with similar anatomy and karyology representing intraspecific variation within *S. lugubris*.

**Family: Dendrocoelidae Hallez, 1892**

**Genus: *Polycladodes* Steinmann, 1910**

**Species: *Polycladodes alba* Steinmann, 1910**

**Material Examined.** H0218, Dretulja Spring, Plaški, Croatia (45.0745° N, 15.3428° E), 13/09/21, coll. Jochen C. Rink, Miquel Vila-Farré, Ludwik Gąsiorowski, Uri Weill and Rick Kluiver, sagittal sections on 20 slides. H0219, Ibid., sagittal sections on 11 slides. H0658, Dretulja Spring, Plaški, Croatia (45.0745° N, 15.3428° E), 19/06/23, coll. Miquel Vila-Farré, sagittal sections on 18 slides.

### Taxonomic discussion

The most distinctive external anatomical feature of *P. alba* is the presence of two separate fields of eyes and its unpigmented colouration (Fig. 2B and Fig. 6A-C). *Polycladodes alba* is further characterised by an anterior intestinal branch that does not reach the level of the eyes (Fig. 6B), a well-developed anterior adhesive organ (Fig. 6B, D), a short pharynx with the mouth located approximately in the middle of the pharyngeal pouch(94) (Fig. 6E), a layer of longitudinal muscles beneath the epidermis of the external pharynx wall(47), predominantly ventral testes (Fig. 6D), a very long penis papilla (Fig. 7A-C), vasa deferentia that open into the anterior section of the seminal vesicle inside the penis papilla (Fig. 7C), and a very long adenodactyl with a very long free papilla (Fig. 7A-B, D).

**Figure 6.**
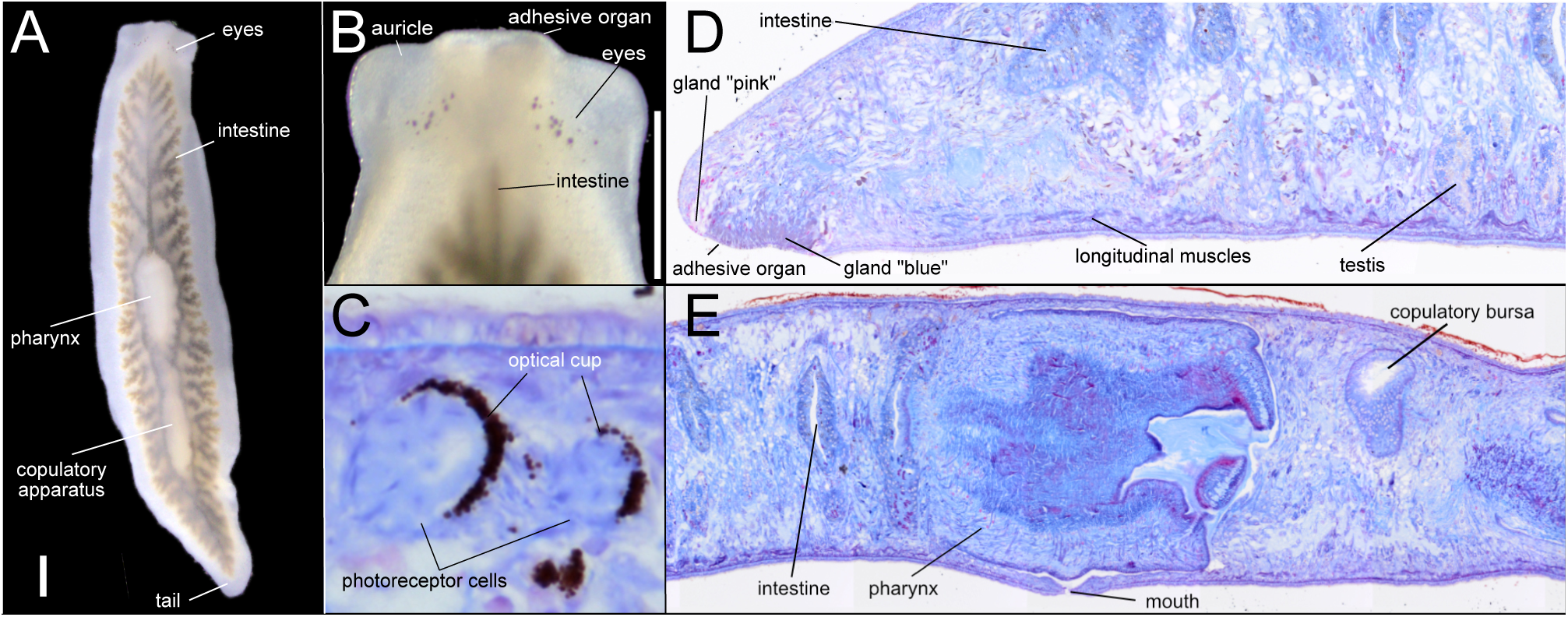
External and internal anatomy of *Polycladodes alba* from Croatia. **A-B.** Live specimens. Scale bar, 1 mm. **A.** Intestinal morphology, revealed by the ingested food. **B**. Detail of the head. **C-D.** Brightfield image of sagittal sections H0218. **C**. Anatomical details of two eyes. **D.** Anterior section of the body showing the adhesive organ. **E.** Middle section of the body showing the central position of the mouth in the pharyngeal pouch.

**Figure 7.**
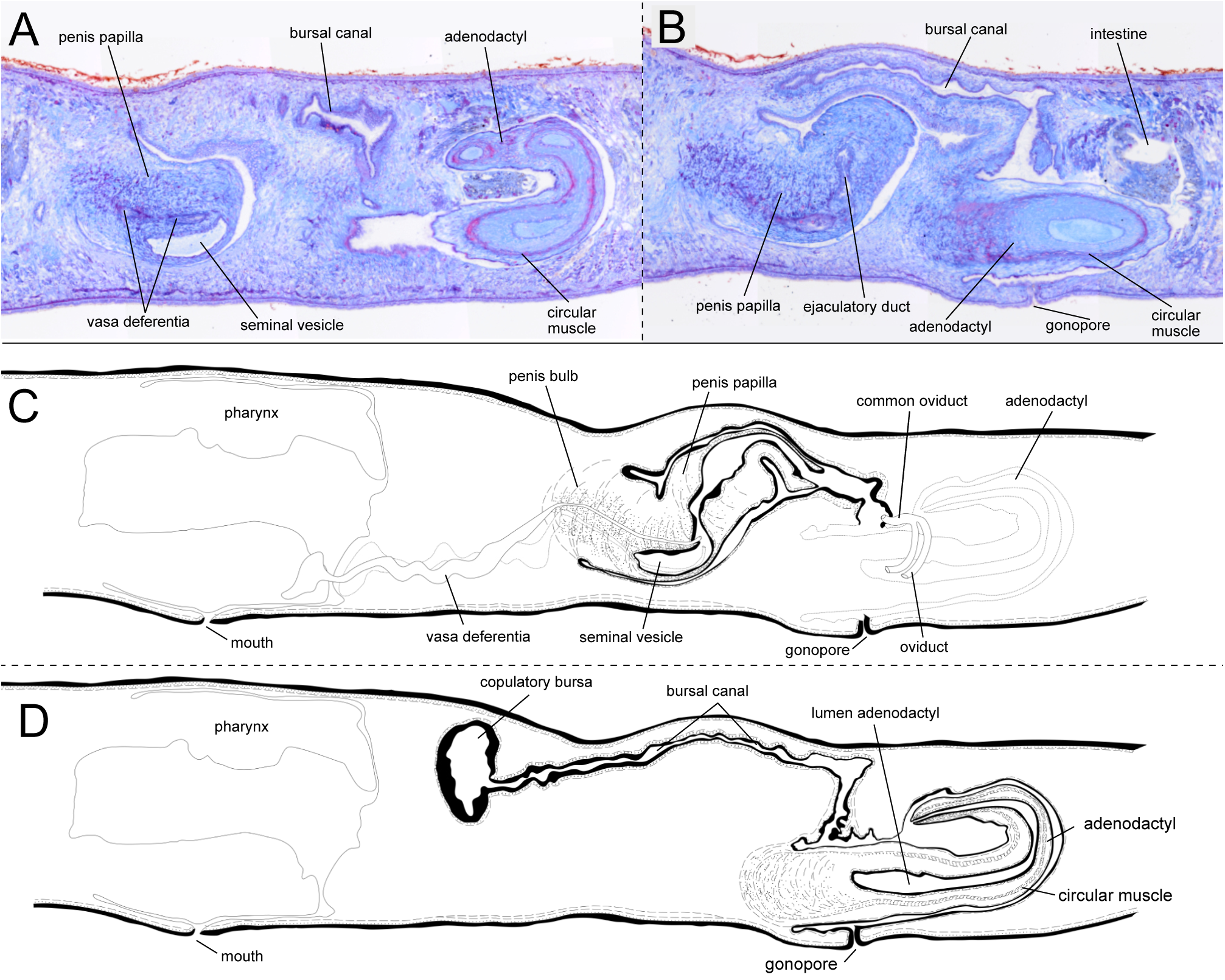
The copulatory apparatus of *Polycladodes alba*. **A-B.** Brightfield image of sagittal sections. H0218. **A.** Detail of the base of the penis papilla and the distal section of the adenodactyl. **B.** Detail of the penis papilla, bursal canal and base of the adenodactyl. **C-D.** Diagrammatic reconstruction of the copulatory apparatus. **C.** Section containing the long penis papilla. **D.** Section containing the copulatory bursa, bursal canal, adenodactyl, and gonopore.

In the Croatian specimens, the testes are predominantly ventral, though some are dorsally located, as observed in individuals H0218 and H0658. The copulatory bursa is small in two analysed specimens (H0218, H0219). The bursal canal is long, consistent with individuals from other populations(95).

Notably, the adhesive organ in the Croatian specimens contains glands. However, it is less developed than that described by de Beauchamp(47), appearing less conical and lacking the “suction cup” shape (Fig. 6D). Of note, the Croatian specimens present in the adenodactyl a circular muscle band with longitudinal muscles ectally and entally distributed to the circular muscle (Fig. 7B, D). An adenodactyl with a reminiscent muscular structure, the “Balkan type of adenodactyl”, is present in several species of *Dendrocoelum* broadly distributed in Europe, including the Western Balkans(96).

**Genus: *Dendrocoelum* O□rsted, 1844**

**Species: *Dendrocoelum pigmentatum* Vila-Farré sp. nov.**

**Material Examined**

Holotype: H0366, Majerovo vrilo, Sinac, Croatia (44.8147° N, 15.3579° E), 13/09/21, coll. Jochen C. Rink, Miquel Vila-Farré, Ludwik Gąsiorowski, Uri Weill and Rick Kluiver, sagittal sections on 8 slides. Paratypes: H0364, ibid., sagittal sections on 11 slides; H0222, ibid., sagittal sections on 13 slides; H0220, ibid., horizontal sections on 9 slides Other material: H0293, Dretulja River, Plaški, Croatia (45.085° N, 15.3663° E), 13/09/21, coll. Jochen C. Rink, Miquel Vila-Farré, Ludwik Gąsiorowski, Uri Weill, and Rick Kluiver, sagittal sections on 16 slides; H0294, ibid. sagittal sections on 33 slides. H0296, Dretulja Spring, Croatia (45.0745° N, 15.3428° E), 13/09/21, coll. Jochen C. Rink, Miquel Vila-Farré, Ludwik Gąsiorowski and Uri Weill, Rick Kluiver, sagittal sections on 14 slides H0297, ibid. sagittal sections on 13 slides. One individual per locality was barcoded; corresponding GenBank accession numbers will be uploaded to GenBank.

### Diagnosis

*Dendrocoelum pigmentatum* **sp. nov.** can be distinguished from its congeners by the presence of body pigmentation, comprising a band along the midline and a transversal band that form a pale dorsal cross-like pattern, a pharynx longer than the adenodactyl, a penis papilla underlined with strong circular muscles, similar or longer than the papilla of the adenodactyl, and by the presence of a spacious cavity coated with a thick layer of muscles before opening into the gonopore.

### Etymology

The epithet is derived from the Latin adjective *pigmentatum*, meaning pigmented, painted, or coloured. It alludes to the presence of body pigmentation in the species, a rare trait in *Dendrocoelum* that is so far only known from species restricted to the Ohrid and Prespa lakes region.

### Habitat

*Dendrocoelum pigmentatum* has been collected from three nearby localities in the Croatian karst (Fig. 8A). One population occurs at Majerovo vrilo (visited by us twice), one of the three major springs of the Gacka River(97), approximately 30 km from the other two and separated by the Mala Kapela mountain range. The Gacka flows 11 km before sinking underground for ∼20 km, re-emerging to ultimately discharge into the Adriatic Sea(98).

**Figure 8.**
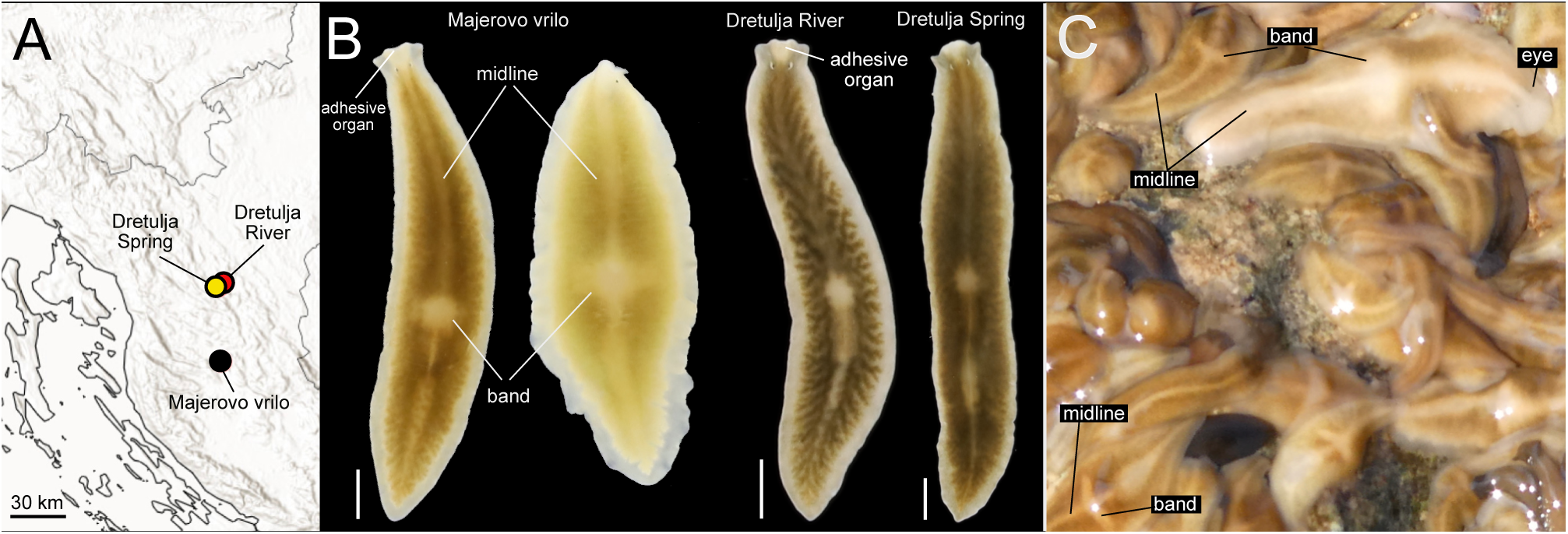
Distribution and pigmentation pattern of *Dendrocoelum pigmentatum* sp. nov. **A.** Distribution of three known populations. **B.** Living animals from the three populations, imaged under laboratory conditions. Animals from Majerovo vrilo were imaged together to show the intrapopulation variability in pigmentation. Scale bar, 2 mm. **C.** Live image of animals in Majerovo vrilo in their habitat. The striking cross-like design formed by the dorsal midline and transverse bands is evident.

The other two populations inhabit the Dretulja River system: one at its source, Dretulja Spring (visited twice), and the other about 2.2 km downstream near Plaški (visited once). The Dretulja is a tributary of the Mrežnica(99), which belongs to the Danube system and ultimately flows into the Black Sea. Despite their proximity, the Majerovo vrilo and Dretulja populations are situated in distinct drainage basins—Adriatic and Black Sea—highlighting potential biogeographic and ecological differentiation.

At Majerovo vrilo (Sinac), the species is abundant in the outlet immediately below the mill. This habitat has a depth of less than 50 cm, a stony bottom, and is interspersed with aquatic vegetation. Here, *D. pigmentatum* co-occurs with *Polycelis felina*, *Polycelis* sp., *Phagocata* sp. and an unidentified white dendrocoelid.

At the outlet of the Dretulja Spring, approximately 7 meters wide, the habitat features a stony substrate and abundant aquatic vegetation. In this locality, *D. pigmentatum* is relatively common and co-occurs with *Polycladodes alba* (rare) and *Polycelis felina* (abundant).

Finally, at the Dretulja River near Plaški, the river measures approximately 10 meters in width and has a depth of around 80 cm. This site is situated upstream of a fish farm. Here, *D. pigmentatum* was less common during our single visit to the locality and coexisted with *Polycelis felina*. In the three localities where it occurs, *D. pigmentatum* is found in flowing water habitats.

### Description

Live specimens with a body length of up to approximately 25 mm and a width of ∼4 mm in the central part of the body and approximately 2.5 mm at the level of the eyes. There is a constriction at the level of the eyes, which are situated apart (Fig. 8B). Anterior end truncated and provided with a pair of lobulated latero-anterior projections and a subterminal anterior adhesive organ (Fig. 8B, 9A). Live animals are pigmented with brown dorsal colouration. In Majerovo vrilo, the degree of pigmentation varies substantially between individuals, with some being much paler than the rest (Fig. 8B, C). In the Dretulja River populations, the observed individuals have a similar brown dorsal colouration (Fig. 8B). The adhesive organ zone has a more creamy pigmentation that forms a triangular area extending backwards to the level of the eyes. A rim in the body margin is lighter or almost unpigmented. There is a pale midline along the anteroposterior axis of the animal, starting behind the eyes and reaching almost the tail. A pale transversal band extends over the region of the pharynx, reaching laterally about halfway between the mid-line and the lateral rim. The pale transversal band and the dorsal midline form a cross-like pattern very conspicuous under field conditions (Fig.8C). In animals imaged under laboratory conditions, this “cross” can resemble a “dot” (Fig. 8B).

**Figure 9.**
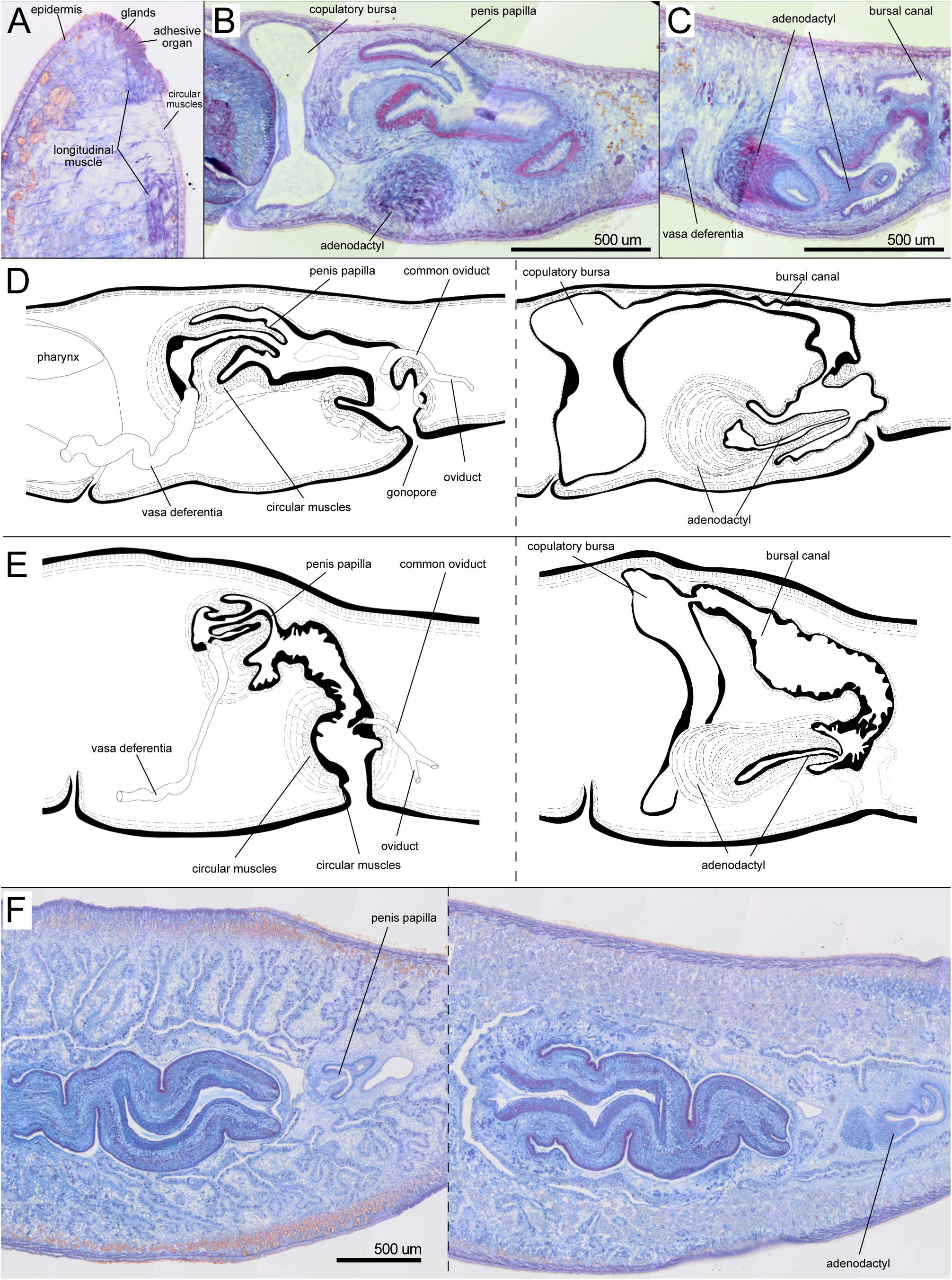
Internal anatomy of *Dendrocoelum pigmentatum* sp. nov. **A** H0222. Brightfield image of sagittal sections. Detail of the adhesive organ**. B-D** H0366 **B-C.** Brightfield image of sagittal sections. B. Detail of the penis papilla and the atrium. **C.** Detail of the adenodactyl. **D.** Diagrammatic reconstruction of the copulatory apparatus. Left, section containing the penis papilla, male atrium and gonopore. Right, section containing the adenodactyl, copulatory bursa and the bursal canal. **E.** H0293. Diagrammatic reconstruction of the copulatory apparatus. Left, section containing the penis papilla, male atrium and gonopore. Right, section containing the adenodactyl, copulatory bursa and the bursal canal. **F.** Brightfield image of horizontal sections H0220. Detail of the pharynx and the copulatory apparatus.

Subterminal anterior adhesive organ is only moderately developed, consisting of a shallow cup made up of epithelial cells, which are pierced by numerous openings of glands (Fig. 9A). The very thick anterior ventral longitudinal body musculature is interrupted at the level of the adhesive organ, in a parenchymal area almost devoid of longitudinal fibres in H0222. Right after, a few longitudinal fibres at the same level as the ventral longitudinal muscles are associated with the adhesive organ (Fig. 9A).

The pharynx is located approximately in the middle of the body and measures less than 1/4th of the body length in specimens from Majerovo vrilo H0222, H0364 and H0366) while it is longer than 1/4th of the body length in specimens from the Dretulja River H0293, H0296). The mouth is situated in a terminal position in the pharyngeal pouch in specimens from Majerovo vrilo H0222, H0364 and H0366) but slightly shifted to the back in the two populations along the Dretulja River H0293, H0294, H0296 and H0297).

The two ovaries occur on the ventral side of the anterior region behind the brain in individuals H0364 and H0366. The oviducts originate from the laterodorsal part of the ovaries and are provided with a slight expansion at their anterior end, the tuba. The oviducts run posteriorly, converging behind the copulatory apparatus to form a single common oviduct (Fig. 9D-E). The common oviduct runs anteriorly and opens into the distal end of the male atrium before it expands to form a broad cavity. The distal part of the oviducts and the first tract of the common oviduct receive the openings of the shell glands in individuals H0366 and H0294.

The numerous well-developed testes can extend from the ventral to the dorsal surface of the body and from behind the ovaries to close to the posterior end of the body. In specimen H0366, the sperm ducts form very large spermiducal vesicles, packed with sperm, between the last section of the pharyngeal pouch and the mouth. The copulatory apparatus is localised behind the pharyngeal pocket. In specimens H0366 and H0293, the copulatory bursa has the shape of a large sack that occupies almost the entire dorso-ventral diameter of the body (Fig. 9B, D, E). The bursa is lined with an epithelium. In H0366, the bursal canal runs posteriorly to the side of the penis and widens at its distal part, forming an enlargement at the end, then turns ventrally and narrows before opening into the atrium of the adenodactyl. In other specimens, the bursal canal widens towards the centre before narrowing when opening on the atrium of the adenodactyl (H0364, H0293).

The penis bulb and the penis papilla are positioned dorsally and laterally to the adenodactyl (Fig. 9D, E, F). The bulb of the penis is anterior to the bulb of the adenodactyl. The muscular penis bulb, formed by longitudinal and circular muscle fibres, is of moderate size. The vasa deferentia penetrate the penis bulb separately at its middle part. Then, they open separately into the lower section of the long seminal vesicle housed by the bulb. The penis papilla is covered by an epithelium at the basal part that becomes thinner at the distal part. The epithelium is underlined by a thick layer of circular muscle in the proximal section of the penis papilla, particularly on the ventral side of the papilla. This thick muscle layer extends in the ventral section of the atrium close to the base of the penis papilla (Fig. 9B, D, E). In specimens H0293 and H0222, the penis papilla appears invaginated into the large papilla lumen (Fig. 9E). The length of the penis papilla (not invaginated) is similar to that of the papilla of the adenodactyl in H0366 (Fig. 9D), and longer in H0296. The penis papilla project into the long and wide male atrium, which is lined by a nucleated epithelium surrounded by a subepithelial layer of circular muscles, followed by a layer of longitudinal fibres. The atrium narrows towards the end before the common oviduct’s opening (Fig. 9D, E). After that point, it forms a spacious cavity coated with a thick layer of circular muscles followed by thick longitudinal muscles in individuals of all the analysed populations (Fig. 9D, E). In individual H0220, the end of the atrium projects into this cavity (Fig. 9F). The cavity opens ventrally in the gonopore.

The very muscular adenodactyl has a ventral and approximately horizontal position. In individual H0366 it consists of a large bulbar part and an elongated papilla (Fig. 9D), similar to specimens H0222 and H0364 from the same population (Majerovo vrilo). Interestingly, in specimens from H0293, H0294 and H0296 (from the Dretulja River and Dretulja Spring), the papilla of the adenodactyl is blunt and short. However, the bulb is also very well developed (Fig. 9E). The papilla is rich in longitudinal and circular muscle fibres. The bulb of the adenodactyl consists of intermingled rows of longitudinal and circular muscle that enter the papilla, which is also muscular. The section of the atrium housing the adenodactyl communicates with the male atrium approximately at the level where the latter expands to form a broad cavity not far from the genital pore.

### Taxonomic discussion

The most characteristic trait of *D. pigmentatum* is the presence of body pigmentation. Therefore, we will restrict our comparative discussion to the pigmented species of the genus *Dendrocoelum*, confined to the Ohrid-Prespa lakes region, that proved phylogenetically close to *D. pigmentatum* in our phylogeny (Fig. 4). The homogeneous anatomy of the copulatory apparatus of those species(85, 100, 101) led Kenk to incorporate other characters to support the species identification, including the pigmentation pattern, as an easily recognisable feature of those species(85), that Stanković considered species-characteristic(102). The dorsal pigment pattern of three pigmented *Dendrocoelum* is reminiscent of that in *D. pigmentatum*: *Dendrocoelum cruciferum* (Stanković, 1969), *Dendrocoelum lacustre* (Stanković, 1938) and *Dendrocoelum lychnidicum* (Stanković, 1969).

*Dendrocoelum cruciferum* is easy to identify by its external morphology(85). The dorsal ground colour is a light yellowish or greyish brown, provided with a broad, lighter rim along the margins. A similar rim is also present in *D. pigmentatum*, although in the latter, the dorsal colouration is probably darker than in *D. cruciferum*, except for some individuals in Majerovo vrilo. In *D. cruciferum*, a distinct, greyish-black stripe runs along the midline of the central area, and a pair of rectangular or rounded dark spots are placed transversely above the region of the pharynx with a crosslike pattern, reminiscent of the crosslike pattern in *D. pigmentatum*. However, although placed in a similar position, in *D. pigmentatum*, the crosslike bar and the midline are lighter than the rest of the dorsal colour and the crosslike band is formed by a single transversal band. Additionally, in *D. cruciferum*, the adhesive organ is highly developed, and the penis is situated at approximately the same level as the adenodactyl(85, 101). In contrast, in *D. pigmentatum*, the adhesive organ is less developed, and the penis is clearly dorsal to the adenodactyl.

The dorsal pattern of *Dendrocoelum lacustre* is characteristic. The animals are poorly pigmented and present scattered streaks of brown pigment along the midline, with two thin dark bands in the lateral edges of the body and two large transverse patches. The size of the animals is up to 1 cm, while *D. pigmentatum* is a much bigger species with markedly different pigmentation. In *D. pigmentatum*, there is a single transversal band and a midline that are paler than the dorsal surface. The band is positioned approximately at the level of the pharynx. Additionally, in *Dendrocoelum lacustre,* the penis is situated slightly dorsal to the adenodactyl, while in *D. pigmentatum*, the penis is dorsal to this structure.

*Dendrocoelum lychnidicum* is a small species up to 6 mm long with a dorsal pigmentation pattern reminiscent of *D. pigmentatum*. In *D. lychnidicum,* the dorsal colouration is of a light chocolate colour that disappears in the head. *D. pigmentatum* is a much longer species, and the head is also pigmented, including the dorsal area around the adhesive organ area. Similar to *D. pigmentatum, D. lychnidicum* presents a light middorsal band and one or more rectangular yellowish rounded spots, forming a crosslike design paler than the ground dorsal pigmentation. Nevertheless, beyond the differences in size and head pigmentation, both species are easy to differentiate by studying their internal anatomy: the length of the pharynx is much shorter than the adenodactyl in *D. lychnidicum,* while in *D. pigmentatum,* it is larger than the adenodactyl. Additionally, in *D. pigmentatum*, the penis papilla is provided with strong circular bands coating the epithelium, and the adenodactyl is proportionally smaller than in *D. lychnidicum*. The seminal vesicle is horizontally oriented in *D. lychnidicum* while vertical in *D. pigmentatum*. Finally, *D. cruciferum*, *D. lacustre* and *D. lychnidicum* inhabit the sublittoral zone of the Ohrid Lake, a different habitat from the river and the karstic springs where *D. pigmentatum* occurs(85, 101).

We previously emphasised the limited availability of barcodes for Dendrocoelids (Fig. 1B) and molecular data in general that we can use to establish comparisons with *D. pigmentatum*. However, the confirmed phylogenetic relationship between *D. pigmentatum* and the pigmented *Dendrocoelum* from Ohrid (Fig. 4) underscores the importance of including pigmented dendrocoelids in comparative analyses. In this context, *D. pigmentatum* is at least distinguishable by barcoding from two pigmented Ohrid forms, *Dendrocoelum* cf. *santinaumi* and *Dendrocoelum maculatum* (Figs. 2b, 4). Furthermore, *D. pigmentatum* individuals from the Dretulja River and Spring cluster together in our barcoding analysis, separate from the individual in Majerovo vrilo (Fig. 2b), supporting the existence of interpopulation genetic variability within the species.

**Family: Planariidae Stimpson, 1857**

**Genus: *Planaria* Mu□ller, 1776**

**Species: *Planaria torva* (Müller, 1774)**

**Material Examined.** H0797, Šarena Jezera, Knin, Croatia (44.02686° N, 16.22285° E), 2013, coll. Jochen C. Rink, sagittal sections on 5 slides. H0798, ibid., sagittal sections on 5 slides. H0799, ibid., sagittal sections on 4 slides. H0800, ibid., sagittal sections on 5 slides. H0801, ibid., sagittal sections on 4 slides.

### Taxonomic discussion

*Planaria torva* is a widely distributed European species, identified in two Croatian localities based on external morphology and barcoding matches to specimens previously studied and catalogued in our planarian collection. Notably, we found no COI accessions for this species in GenBank, emphasising the need to document its characteristics. The Croatian specimens conform to the species’ known external appearance: they measure up to ∼1.2 cm in length, have a truncated head with two eyes (Fig. 10A), and exhibit a uniform brown dorsal colouration. This description broadly matches descriptions of *P. torva* from the United Kingdom (83) and Herzegovina(84).

**Figure 10.**
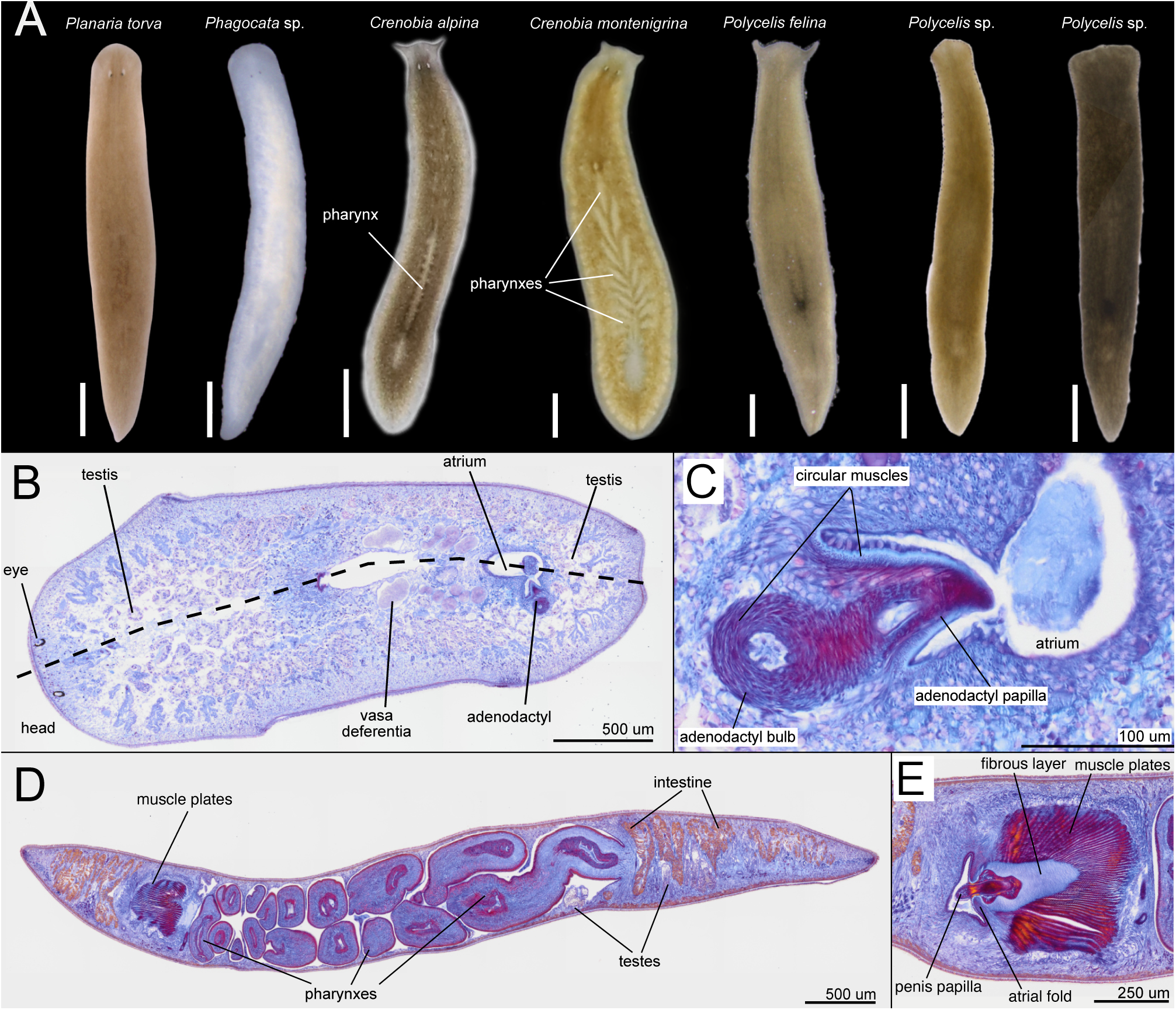
Diversity of planariids. **A.** Living planarian specimens representing the planariids collected during our expeditions. Scale bar, 1 mm. **B-C.** *Planaria torva*. H0798 Brightfield image of horizontal sections. **B.** Testis distribution and adenodactyl position. **C.** Anatomical detail of the adenodactyl. **D-E.** *Crenobia montenigrina*. H0659. Brightfield image of sagittal sections. **D.** Longitudinal section showing the presence of multiple pharynxes, the position of the testes, and the muscle plates of the copulatory apparatus. **E.** Copulatory apparatus.

Anatomically, *P. torva* is distinguished among European planariids by the presence of an adenodactyl—a muscular organ projecting into the atrium. In the Croatian specimens, this organ is visible in histological sections (Fig. 10B-C). This trait, coupled with the barcoding results, confirms the taxonomic designation of these specimens as *Planaria torva*.

**Genus: *Crenobia* Kenk, 1930**

**Species: *Crenobia montenigrina* (Mrázek, 1904)**

**Material Examined.** H0659, Vuković vrilo, Civljane, Croatia (43.9654° N, 16.4129° E), 2023, coll. Miquel Vila-Farré, sagittal sections on 10 slides.

### Taxonomic discussion

Sluys[44] recently reevaluated the morphological variability within the genus *Crenobia*. Guided by these findings, we analysed the Croatian *Crenobia montenigrina* specimens to validate their taxonomic assignment. Those specimens exhibit the expected polypharyngeal trait (Fig. 10A, D); the anteriorly shifted mouth opening; large, primarily ventral testes extending posteriorly beyond the root of the first pharynx (Fig. 10D); a pronounced curvature of the sperm ducts; and a thick muscular coat surrounding the atrium (muscle plates) and a fibrous plate characteristic of the genus *Crenobia* (Fig. 10E). In summary, the anatomy of the Croatian specimens is consistent with that of *C. montenigrina*(51).

### General discussion

Our study’s results underscore the importance of integrative approaches in documenting and ultimately preserving biodiversity. The inconclusive barcoding results for specific taxa highlight the limitations of relying solely on COI sequences for taxonomic resolution, particularly when the reference library suffers from taxonomic biases. Notably, COI sequences for common and abundant European freshwater planarians with potentially relevant environmental roles, such as *Dendrocoelum lacteum*, *Planaria torva*, *Polycelis nigra* or *Polycelis tenuis,* are either poorly represented or absent in the current GenBank record. Notably, the abundance of representatives from some of these groups in Croatia prompted us to generate new sequences now incorporated into public repositories. These sequences, and additional ones produced in this study from other geographic areas from all over Europe, increase substantially the number of available COI sequences for European planariids and dendrocoelids, e.g., for *D. lacteum*, contributing to the goal of a taxonomically curated barcoding reference library for planarians.

Our integrative approach further provided new insights into the taxonomy of several taxa. Our investigation of *Schmidtea lugubris* revealed the existence of two genetically distinct lineages, the Central and Adriatic haplotype clades, which were morphologically and karyologically indistinguishable. Even in the case that a more detailed analysis of the two clades should uncover clade-specific differences, our transcriptomic branch-length analysis suggests that the genetic difference between the lineages remains below the level observed between established sister species within *Schmidtea*. This supports the interpretation that the two clades represent intraspecific genetic diversity rather than distinct species.

The geographic distribution of the two major genetic clades in *S. lugubris* is compatible with a hypothetical origin from the southern glacial refugia, particularly for the Adriatic clade. Species of Mediterranean origin, isolated in southern European peninsulas during glacial periods, frequently give rise to differentiated genetic lineages. These lineages may remain confined to their respective peninsulas (Balkans, Italy, and Iberia) or expand via diverse dispersal routes(103–107). This pattern is observed in the Balkans across various organisms(107–110), including aquatic species(111–114). *S. lugubris* could represent an additional case of a species clade surviving glacial periods in the Balkans, the Adriatic clade. However, due to our limited sampling in much of the Western and Southern Balkans, the precise geographic extent of the Adriatic clade remains uncertain. Further research in the Balkans, Pannonian Basin, and surrounding regions is essential to resolve the species’ phylogeography. The presence of highly similar haplotypes in the Central clade across its northern range—from Italy and the United Kingdom to southern Sweden—is compatible with a postglacial expansion from a southern refugium. Additionally, the genetic diversity observed in the eastern part of the Central clade and within the Adriatic clade indicates a complex biogeographic history that warrants further investigation. The biogeography of European planarians inhabiting lowlands (e.g., *S. polychroa*, *D. lacteum*, *P. tenuis* or *S. lugubris*) is barely studied(115), in contrast with cold-adapted species like *Crenobia alpina*(39, 116) or the *Dugesia* in the Mediterranean peninsulas(117–119). Our new primers, COI reference sequences and transcriptomes offer an entry point for addressing this gap. Finally, understanding the genetic similarities and differences between the two *S. lugubris* clades deserves further investigation. Modelling the geographic distribution of both clades using species distribution modelling (SDM) software based on occurrence records(118, 120) can potentially identify regions where both clades occur in proximity, which will help clarify the phylogeography and evolutionary history of this recent divergence, e.g., if both clades are in contact or isolated. At the same time, genomic resources for *S. lugubris*(74) and transcriptomes for individuals of both clades can facilitate understanding the genomic differences between the Adriatic and the Central clades (e.g., synteny analysis). Therefore, the complex population history of *S. lugubris* presents an opportunity to explore patterns of planarian genome evolution at the species level.

*Polycladodes alba* is a widespread but rarely studied Centro European species (see summary in(48))(121, 122)). Komárek(43) described the cave species *Sorocelopsis decemoculata* Komárek 1919 based on a single specimen from the cave system Đulin ponor – Medvedica near Ogulin, Croatia. Later, De Beauchamp(47) considered this record an individual of *P. alba.* If *S. decemoculata* is a synonym of *P. alba*, our finding and systematic description of the collected specimens in Croatia mark the species’ reappearance in the country after over 100 years.

The discovery of a new pigmented *Dendrocoelum* in the Northern Dinaric Karst, far from Lake Ohrid, is striking. Although we cannot rule out a human introduction of planarians(123, 124), the presence of significant genetic differences (COI barcoding tree) and morphological variation (e.g., the dorsal pigmentation pattern and the anatomy of the adenodactyl) between populations suggest otherwise. The two populations of *<u>D. pigmentatum</u>* in the Dretulja River are separated from those at Majerovo vrilo by a mountain range and belong to two different major drainage systems, the Black Sea and Adriatic basins (see the description of the species for details).

All the pigmented and a few unpigmented forms of *Dendrocoelum* were formerly assigned to *Neodendrocoelum*, a clade of discussed taxonomic validity (85, 125). Interestingly, the unpigmented species have been recorded in localities distant from the Ohrid, including Croatia (*Dendrocoelum subterraneum*(100) and, with no taxonomical evidence, *Dendrocoelum plesiophthalmum*(52)), Herzegovina (*Dendrocoelum plesiophthalmum*), or in the Western and Southern Balkans (*Dendrocoelum nausicaae*)(85, 102). Our phylogenetic analysis supports a close relationship between the pigmented *D. pigmentatum* and the pigmented *Dendrocoelum* in the Ohrid area, and it is compatible with the existence of a phylogenetic clade of *Dendrocoelum* species (pigmented and unpigmented) that is widely distributed in the Western Balkans and includes the occurrence of pigmented species further north than previously suspected. Hence, our data add an intriguing dimension to the long-debated phylogenetic affinities of these peculiar animals(47, 85, 102, 126), which has proven difficult to resolve by morphological analysis alone(85). The transcriptomes generated in this study, combined with the description of a new pigmented *Dendrocoelum* from Croatia, provide a valuable entry point for addressing these long-standing questions from a molecular perspective. Additionally, our taxonomic description of the unusually large and charismatic (by planarian standards) *Dendrocoelum pigmentatum* can help in conservation efforts for karstic springs in Croatia.

Our sampling in Croatia was limited in scope and restricted to surface waters. Nevertheless, it increased the number of known freshwater planarian species in Croatia from eight to sixteen, with about 35 new planarian records (Fig. 11 A, B). While some taxonomically curated records published may have been missed in our bibliographic research, most species cited in the literature are reported from only one or a few localities, making our contribution of multiple sites with planarians more relevant. Given our limited sampling effort, additional sampling in surface waters and caves will likely raise the species diversity in the area.

**Figure 11.**
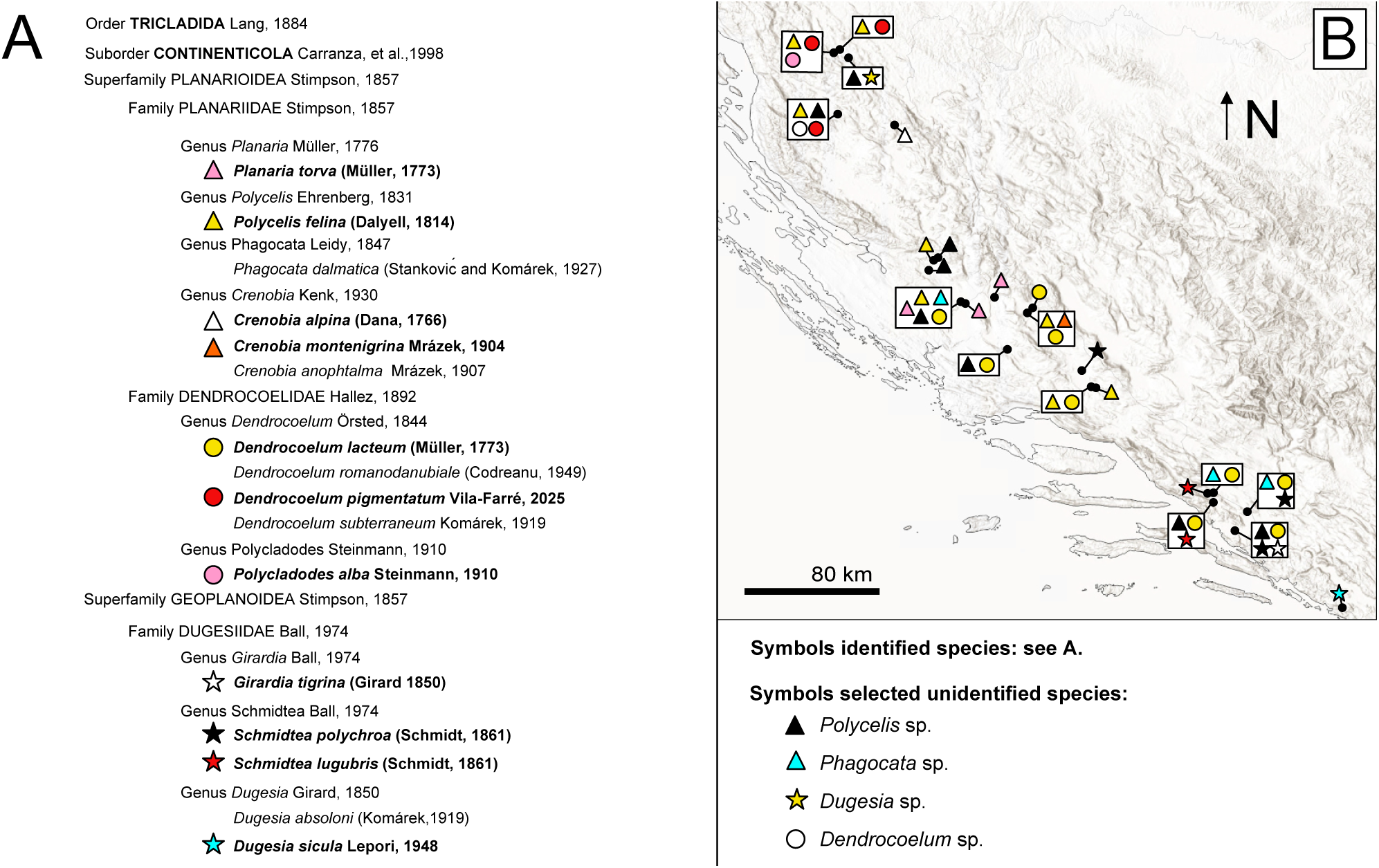
A. Updated list of Croatian Continenticola, excluding terrestrial species. Bold: species studied in this paper. To the left of each species, symbol used in **B** to identify our collection sites for the species. **B.** Distributional records of the species investigated in this study in our collection sites, and of selected samples identified to the genus level.

### Conclusions

Further efforts are needed to expand molecular and morphological datasets for planarians, particularly for rare and understudied species. Adding multiple planarian COI barcodes from understudied groups represents an entry point for the future generation of a geographically broad and taxonomically inclusive COI barcoding reference library for freshwater planarians. Additionally, discovering differentiated genetic lineages of *Schmidtea lugubris*, identifying taxonomic ambiguities, and describing new species underscores the need for further research in species-rich areas like Southeastern Europe. While areas such as the Peloponnese, southern Ionian islands, the Aegean islands, the Iberian Peninsula or Sardinia have recently been investigated using molecular tools(117, 119, 127, 128), others, such as the south part of the Dinarides and the Albanids, remain largely unexplored using molecular techniques. Ultimately, this work lays the foundation for a more comprehensive framework for the taxonomy and phylogenetics of European freshwater planarians, particularly in the Western Balkans, and contributes to broader efforts to document and thus conserve aquatic biodiversity.

## Declarations

### Ethics approval and consent to participate

Not applicable.

### Consent for publication

Not applicable.

### Availability of data and materials

All newly reported gene sequences, new raw RNA sequencing, new transcriptome assemblies and histological slides will be submitted to public repositories.

### Authors’ contributions

M.V.-F. and J.C.R. conceptualised the study; M.A.G. and M.V.-F. designed the primers; M.V.-F., J.C.R., U.W, R.K., L.G., L.K., H.B. collected samples; M.V.-F., M.B., F.F.-S., T.B., performed the experiments and optimised protocols; F. F-S., M.V-F. and M.R. generated and curated the DNA sequences; M.V.-F. and M. B. performed anatomical reconstructions; M.V.F. developed the taxonomic evaluation of the available material; J.B. performed the data analysis for barcoding and phylogeny; M.V.-F. and J.C.R. wrote the manuscript and revised the manuscript text.

## Acknowledgements

We thank Sajmir Beqiraj, Maria Qarri, Marko Lukić, Lada Jovović, and Magdalena Grgić for fieldwork support and Delia Niehaus for support with experimental work. We also thank J. Krull and the MPI-NAT animal services staff for worm care and Rink laboratory members for lively discussions.

---

**Additional File 1. Primers used for amplification and sequencing.**

**Additional File 2. Primer and COI gene sequences used in this study**. GenBank accession IDs are listed for all the sequences. Note that only the primer sequences for new sequences are listed. FJ646985 and FJ646940: sequence generated from two non-overlapping GenBank COI sequences (FJ646985 and FJ646940).

**Additional File 3. Haplotype network for the mitochondrial gene COI I from Dataset 2 (sequence length 702 bp).** Each circle represents a different haplotype, and the size of the circle indicates the frequency of each haplotype. Black bars represent intermediate (non-present) haplotypes, and lines connecting haplotypes (existing or not) represent one nucleotide change. The colour scheme differentiates the major haplotype clades (see legend). The use of a dataset with long sequences highlights the existence of notable genetic differences within the Adriatic clade.

**Additional File 4. The copulatory apparatus of *S. lugubris* is broadly similar in British and Croatian specimens. A-B.** Specimen H0273 from Croatia. Scale bar, 500 µm**. A.** Diagrammatic reconstruction of the copulatory apparatus. **B.** Brightfield image of sagittal sections shows the nipple and the muscular bulb. **C-D.** Specimen H0803 from United Kingdom. Scale bar, 500 µm **C.** Diagrammatic reconstruction of the copulatory apparatus. **D.** Brightfield image of sagittal shows the nipple and the muscular bulb.

**Additional File 5. Comparison of the anatomy of *S. lugubris* in individuals of the Adriatic and Central clades. A.** Diagrammatic reconstruction of the copulatory apparatus highlighting the anatomical areas of interest. **B-C.** Brightfield image of sagittal sections. Scale bar, µm. **B.** Specimens H0273, H0275 from Croatian (upper) and H0814 and H0813 from Montenegro (bottom). **C.** Specimens H0804, H0803 and H0802 from United Kingdom (upper) and H0805, H0807, and H0806 from Italy (bottom).

