## Supplementary figures and images for "An integrative taxonomic approach reveals unexplored diversity in Croatian planarians"

### Additional File 3

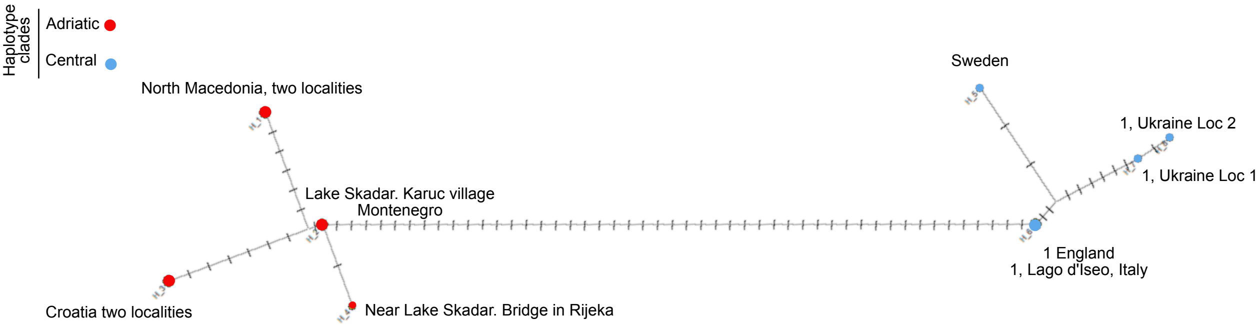

### Additional File 4

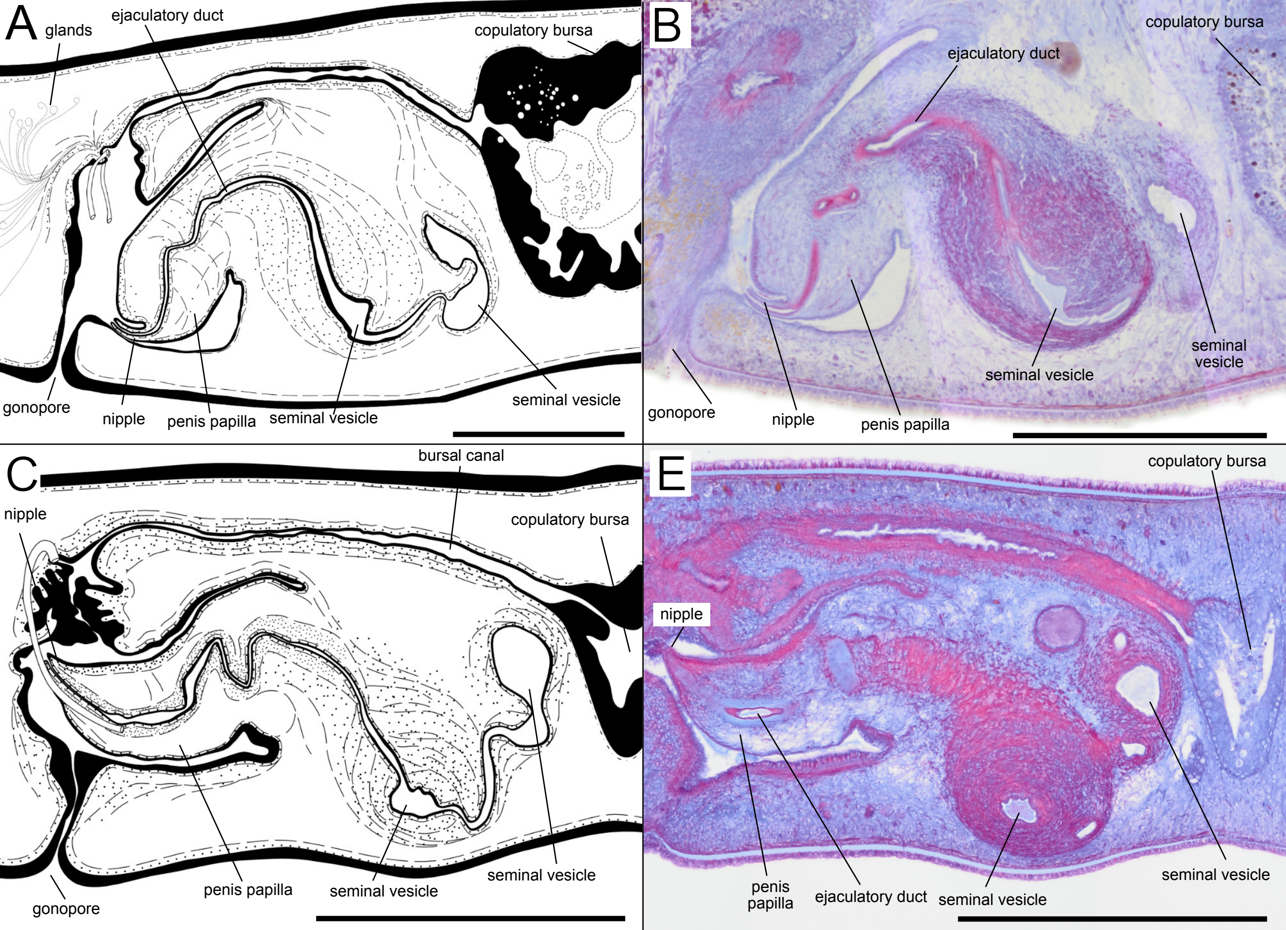

### Additional File 5

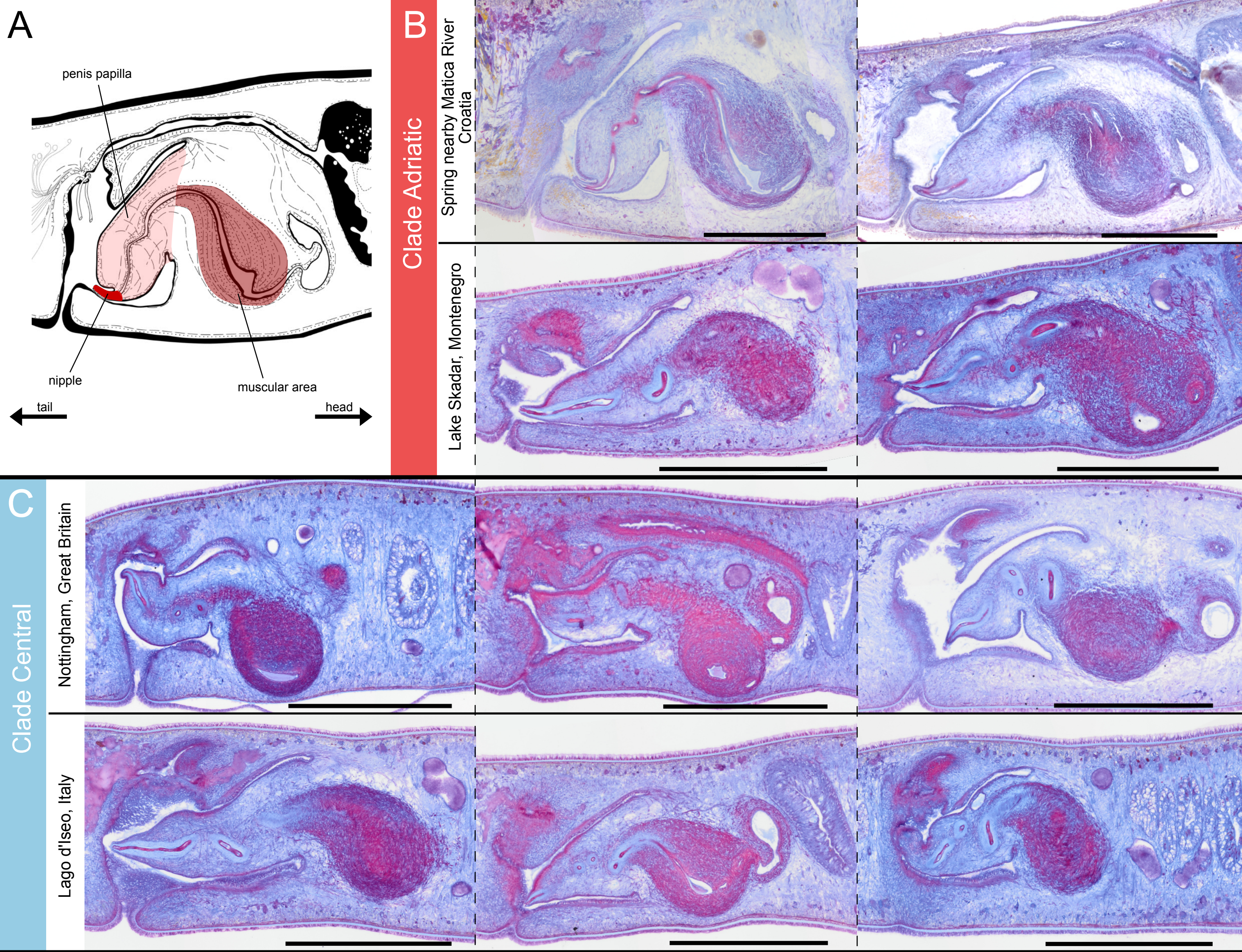
